# Targeting Diffuse Midline Glioma with a novel anti-CD99 Antibody

**DOI:** 10.1101/2024.03.19.585814

**Authors:** Ilango Balakrishnan, Krishna Madhavan, Angela M Pierce, Joshua Michlin, Breauna Brunt, Senthilnath Lakshmana Chetty, Dong Wang, John DeSisto, Zachary Nuss, Nathan Davidson, Andrew Donson, Kenneth Jones, Siddhartha Mitra, Adam Green, Nathan Dahl, Rajeev Vibhakar, Sujatha Venkataraman

**Author notes:** These authors contributed equally (Ilango Balakrishnan, Krishna Madhavan). Co-corresponding and senior authors Correspondence: Sujatha Venkataraman, Department of Pediatrics, University of Colorado Denver, Aurora, CO, 80045, USA. Rajeev Vibhakar, Department of Pediatrics, University of Colorado Denver, Aurora, CO, 80045, USA.

## Abstract

Diffuse midline gliomas (DMGs) are devastating brain tumors that occur primarily in children. The salient feature of these tumors is the presence of a H3K27M mutation (K27M), associated with the worst prognosis. We identified the cell surface antigen CD99 as notably expressed in DMGs, particularly in K27M^+^DMGs. We found that the increased expression of CD99 in K27M^+^DMGs was a result of the onco-histone K27M mutation. In K27M^+^DMG cells, CD99 inactivation impaired tumor growth by inducing cell differentiation, indicating an oncogenic role of CD99 enabled by blocking differentiation. We then developed a novel therapeutic anti-CD99 chimeric antibody, 10D1, with a membrane-proximal binding epitope, and evaluated its antitumor efficacy in preclinical models of K27M^+^DMG. 10D1 suppressed DMG growth *in vitro* and *in vivo* by inducing apoptosis. When combined with radiation treatment, 10D1 exhibited improved antitumor efficiency and xenograft survival, providing a strong justification for its clinical development as a therapy for DMGs.

**Statement of Significance:** This study emphasizes that CD99 overexpression occurs due to the H3K27M mutation in Diffuse Midline Gliomas (DMGs). This heightened expression suppresses apoptosis, inhibits differentiation, and induces radio-resistance in DMGs. This research justifies using a novel CD99 antibody alone or combined with radiation therapy in human pediatric clinical trials.

## Introduction

Diffuse midline gliomas (DMGs) are devastating brain tumors that occur in children (1). These tumors arise from midline brain structures, including the thalamus, pons, and spinal cord, and tend to infiltrate the surrounding brain tissue, making it challenging to remove surgically (2). Over the past four decades, the prognosis has remained equally dismal, with less than 5% of patients alive two years after diagnosis (3,4). Radiotherapy is the standard treatment for DMG and often results in temporary tumor reduction and clinical improvement (2,3). Combination chemotherapy and unselected single-agent targeted therapy trials have not improved the prognosis (3–5). The breakthrough came when pontine biopsies were deemed safe, and tissue became available for the analysis of tumor material (6). This led to the discovery of characteristic mutations in the *H3.3, H3F3A,* or *HIST1H3B* histone genes (7,8). These recurrent histone mutations result in a Lys27Met (K27M) substitution and are found in 74–85% of all WHO grade 4 DMG tumors (9).

Functional studies have suggested that the H3K27M mutation leads to the inactivation of polycomb repressive complex 2 (PRC2) through the sequestration of EZH2(10–12). This results in the global hypomethylation of K27Me3-bound promoters, with consequent transcriptional de-repression of these loci (12). K27M is also highly associated with transcriptionally active chromatin, mediated by EZH2 and BRD4 (13,14). Engineered neural progenitor cells (NPCs) expressing the K27M mutation maintain a stem-like undifferentiated gene signature (15). Furthermore, the overexpression of the K27M transgene cooperates with other factors, including TP53 and PDGFR, to drive DMG tumorigenesis in murine models (10,16). While these studies give insight into driver mutations, they have not yet resulted in new therapeutic options (8,17,18).

To address the problem of identifying therapeutic targets, we examined specific gene expression changes mediated by the K27M mutation. We identified MIC2 as a gene tightly regulated by the H3K27M mutation. *MIC2* encodes CD99, a cell surface glycoprotein (19). CD99 has been shown to enhance cell migration, induce tumor metastasis, and inhibit differentiation in several oncogenic states, including Ewing Sarcoma and acute myeloid leukemia (20–22). Here, we show that the K27M mutation regulates CD99 at the genomic level and is critical for K27M^+^DMG cell viability. In K27M^+^DMGs, genetic depletion of CD99 results in decreased stem cell renewal and increased stem cell differentiation. Mechanistically, we demonstrated that CD99 engages in an anti-apoptotic pathway via integrin-linked kinase (ILK1). We developed a novel IgG4-based recombinant antibody targeting CD99. This antibody blocks CD99-mediated intracellular signaling, resulting in cell death.

*In vivo*, CD99 blockade decreased tumor DMG growth and increased the survival of orthotopic xenograft tumor-bearing mice. Radiation treatment is the only current standard of care for DMG. Importantly, DMG cells subjected to fractionated radiation enhanced the expression of CD99 in these cells. Subsequently, CD99 blockade combined with radiation treatment resulted in significant tumor regression and long-term survival of tumor-bearing mice. We demonstrated a novel mechanism that regulates K27M cell fitness and established a new antibody as a therapeutic agent for this devastating tumor.

## Results

### The oncogenic K27M mutation increases CD99 expression in histone H3 wildtype neoplastic cells

To understand the biological effect of the oncogenic H3.3K27M (K27M) mutation in DMGs, pediatric patient-derived HSJD-GBM01 cells that harbor wildtype histone H3.3 was transduced with the K27M mutant transgene (**Supplementary Figure S1A(i)**) and transcriptomic analysis of the paired isogenic samples was performed. One of the genes uniquely upregulated by expression of K27M mutation was CD99/*MIC2* (**Figure 1A, and Table S4**). CD99 is a cell surface glycoprotein encoded by the *MIC2* gene previously implicated in multiple tumor types, including AML (23) and EWS (21,24,25). Both mRNA and protein analyses confirmed that the increase in CD99 was a result of the overexpression of the K27M mutation in GBM01 cells (**Figure 1B(i), Supplementary Figures S1B(i), C(i)**). Subsequently, overexpression of the K27M mutant transgene in a WT DMG cell line (VUMC-DIPG10) resulted in a similar increase in CD99 mRNA and protein expression (**Figure 1B(ii)**, **Supplementary Figures S1A(ii), B(ii), C(ii)**). Interestingly, analogous K27M overexpression in fetal-derived normal human astrocyte (NHA) cells showed no significant change in CD99, suggesting that the simple genetic overexpression of the oncohistone K27M alone is not sufficient in normal cells and requires a transformed phenotype to alter CD99 expression (**Figure 1B(iii), Supplementary Figures S1A(iii), B(iii)**). Conversely, CRISPR/cas9-mediated deletion of the K27M mutation from K27M^+^-BT245 DMG cells (26) resulted in a loss of CD99 (**Figure 1C**, **Supplementary Figure S1D(i-iii)**). It showed a similar loss in the expression of another cell surface antigen, GD2, as measured by flow cytometry (**Supplementary Figure S1D(iv)**). To further explore these changes in CD99 within the context of the K27M mutation, histone 3 WT (H3WT) transgenes were overexpressed in K27M^+^HSJD-DIPG007 cells and paired isogenic cell lines were analyzed. The overexpression of H3WT was confirmed by the increase in K27Me3 histone protein, and these cells showed a significant decrease in CD99 at both the mRNA and protein levels (**Supplementary Figure S1E**).

**Figure 1:**
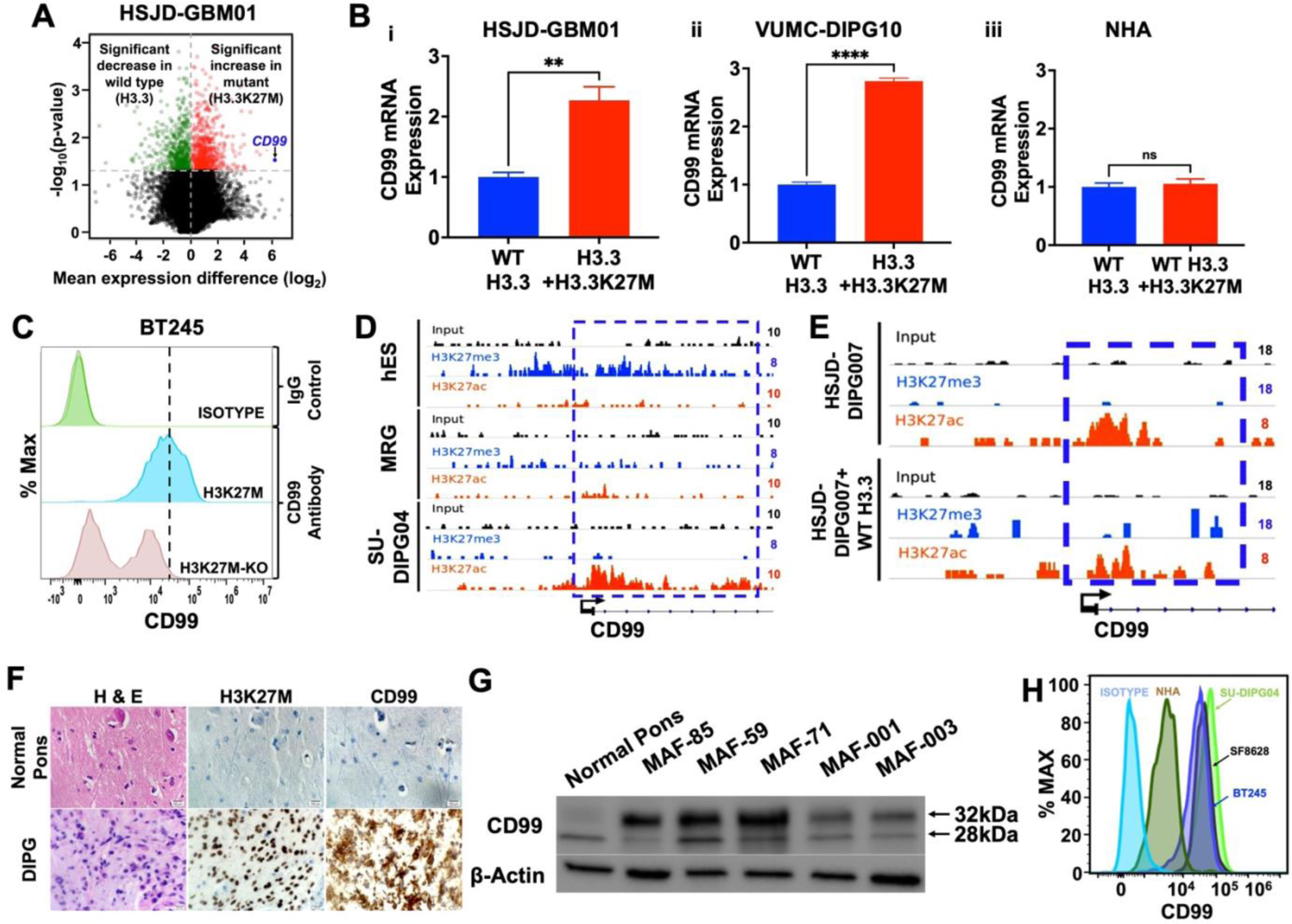
The increase in CD99 expression is associated with H3K27M mutation in neoplastic cells. A. Volcano plot showing the statistical significance as -log_10_(p-value) plotted against the log_2_(fold change) between the genes expressed by HSJD-GBM01 transduced with K27M mutant at the H3.3 locus and the unmodified HSJD-GBM01 (control) cells. HSJD-GBM01 transduced with K27M mutant demonstrated a several-fold increase in CD99 (indicated with blue arrow) compared to the control cells. B. CD99 mRNA expression in the following paired cell lines transduced with H3.3K27M: (i) HSJD-GBM01, (ii) VUMC-DIPG10, and (iii) normal human astrocytes (NHA). (ns = not significant, **p<0.01, ****p<0.0001) C. Flow plot showing a significant decrease in the expression of CD99 in K27M mutant deleted BT245 cells as compared to the parental BT245 cells. D. ChIP-seq signal peaks of H3K27Me3 and H3K27ac in human embryonic stem cells (hES), mid-radial glia (MRG), and DMG (SU-DIPG04) cells suggesting an increase in the activating H3K27ac at the CD99 promoter region in the DMG cells but not in the hES or MRG cells. E. ChIP-seq signal peaks of the paired samples of HSJD-DIPG007 and cells transduced with the H3.3 wildtype transgene reveal a decrease in the acetylation (H3K27ac) activating mark and an increase in trimethylation (H3K27me3) repressive marker at the CD99 promoter region in compared to K27M Mutant parental HSJD-DIPG007 cells. F. Representative immunohistochemical (IHC) staining showing expression of CD99 in K27M-DMG tumor and its absence in normal pons. K27M staining was used to confirm the mutation status of the tumor. G. CD99 is highly expressed in DMG patient tumors as compared to normal pons by western blotting. The molecular weight of the long (32kDa) and short (28kDa) isoforms are denoted next to the western blot image. H. Flow plot showing the higher expression of CD99 in DMG cell lines in comparison to NHA. Data shown as the mean ± SEM. (See also **Supplementary Figures S1 and S2 and Table S4**)

Next, we investigated whether these transcriptional and translational changes in CD99 after K27M modification in neoplastic cells correlated with chromatin activation. Initially, ChIP-sequencing (ChIP-seq) data on DMG and normal samples that were publicly available (27) were reanalyzed for the expression of histone-activating and repressive marks at the CD99 promoter region. From this analysis, at the promoter region of CD99, high levels of activating H3K27Ac marks in K27M^+^SU-DIPG04 cells and increased levels of repressive K27Me3 marks in normal mid-radial glial (MRG) and human embryonic stem cells (hES) were observed (**Figure 1D**). We then performed ChIP-seq using isogenic paired cell lines, K27M^+^HSJD-DIPG007, that were transduced with the H3.3WT transgene. Analysis revealed a decrease in H3K27Ac activation marks and a moderate increase in repressive K27Me3 marks at the CD99 promoter when compared to its parent isogenic cell line (**Figure 1E**). These data indicate that oncohistones control changes in CD99 expression at the genomic level. Together, these results suggest that the increased expression of CD99 in K27M^+^DMGs is a product of the oncogenic K27M mutation that is regulated at the genomic level.

### CD99 is highly expressed in DMGs

Initial analysis of CD99 gene expression in the DMG patient samples (N=36, pediatric DMGs) from the publicly available R2 database revealed that its expression is significantly higher in DMGs than in normal brain regions (consisting of both adult and pediatric tissues) (**Supplementary Figure S2A**). Subsequently, Immunohistochemical (IHC) analysis of a cohort of K27M^+^DMG patient samples and normal pons collected at our center demonstrated that CD99 expression was higher in K27M^+^DMG than in WT DMG patient tumors, whereas it was absent in the normal pons, as shown in **Figure 1F** and **Supplementary Figure S2B.** CD99 is expressed as two major distinct isoforms because of alternate splicing of the encoding gene, MIC2, a long-form containing 185 amino acids (LF, 32kDa,) which has been previously shown as the active form of CD99 and a short form containing 161 amino acid that harbors deletion of the cytoplasmic region (SF, 28 kDa) (28). To identify the preferential expression of the two isoforms in DMG, CD99 protein expression was examined by western blotting in K27M^+^DMG patient samples. Interestingly, as depicted in **Figure 1G**, uniform overexpression of the long active form was observed compared to that of its short form in primary patient tumors and all DMG cell lines tested (**Supplementary Figure S2C**). On the other hand, normal pons predominantly expressed the short form and not the long form of CD99. Based on these data we propose that the K27M mutation preferentially induces the long active form of CD99. To test this effect of K27M mutation on the CD99 long isoform, we selected the ONS76, a medulloblastoma cell line, to transduce with the K27M transgene as these neoplastic cells express H3 WT and low levels of CD99 (**Supplementary Figure S2D, E**). Overexpression of the K27M mutation in ONS76 significantly augmented the expression of CD99 and specifically increased the expression of the long form of CD99, but not the short form (**Supplementary Figures S2F, G**). This suggests a possible selective dependency of K27M^+^DMG tumor cells on the active form of CD99 to enhance proliferation. Overexpression of the long form of CD99 in ONS76 cells led to an increase in cell proliferation, whereas no such increase was observed with overexpression of the short form of CD99 (**Supplementary Figures S2H, I**). Our findings suggest that the long form of CD99 may play a more significant role in cell proliferation than the short form.

By examining the cell surface expression of CD99 in DMG cell lines by flow cytometry, using an antibody specific for the CD99 long form (0662), it was found that (1) all K27M^+^DMG cell lines expressed high levels of CD99 compared to the fetal NHA (**Figure 1H**) and (2) while both K27M^+^ and histone WT DMG cells express CD99 on the cell surface, its expression is higher in the K27M^+^DMGs compared to that of the H3 WT DMG cells (**Supplementary Figure S2J**), further demonstrating that the oncohistone regulates the increased expression of CD99 in K27M^+^DMGs. In addition, single-cell RNA sequencing of a small cohort of K27M^+^DMG patient tumors collected at our center (n=4) demonstrated a uniformly high expression of CD99 in K27M^+^ tumor cells (**Supplementary Figure S2K**).

CD99 has been reported to be expressed in other cells. Using tissue arrays, we found that the spleen and thymus exhibited an abundance of CD99, as previously reported. In contrast, many normal brain tissues showed minimal or no CD99 expression (**Supplementary Figure S2L).**

### CD99 knockdown inhibits DMG cell proliferation

To understand the functional role of CD99 in DMGs, the expression of CD99 was depleted in K27M^+^DMG cell lines using lentiviral shRNAs (**Supplementary Figure S3A, B**). Using these modified cell lines, the potential effects of CD99 on cell proliferation were examined by xCELLigence and incucyte. The results demonstrated that DMG cells transduced with shCD99 showed a significant decrease in proliferation compared to shNull-transduced cells (**Figure 2A**, **Supplementary Figure S3C**).

**Figure 2:**
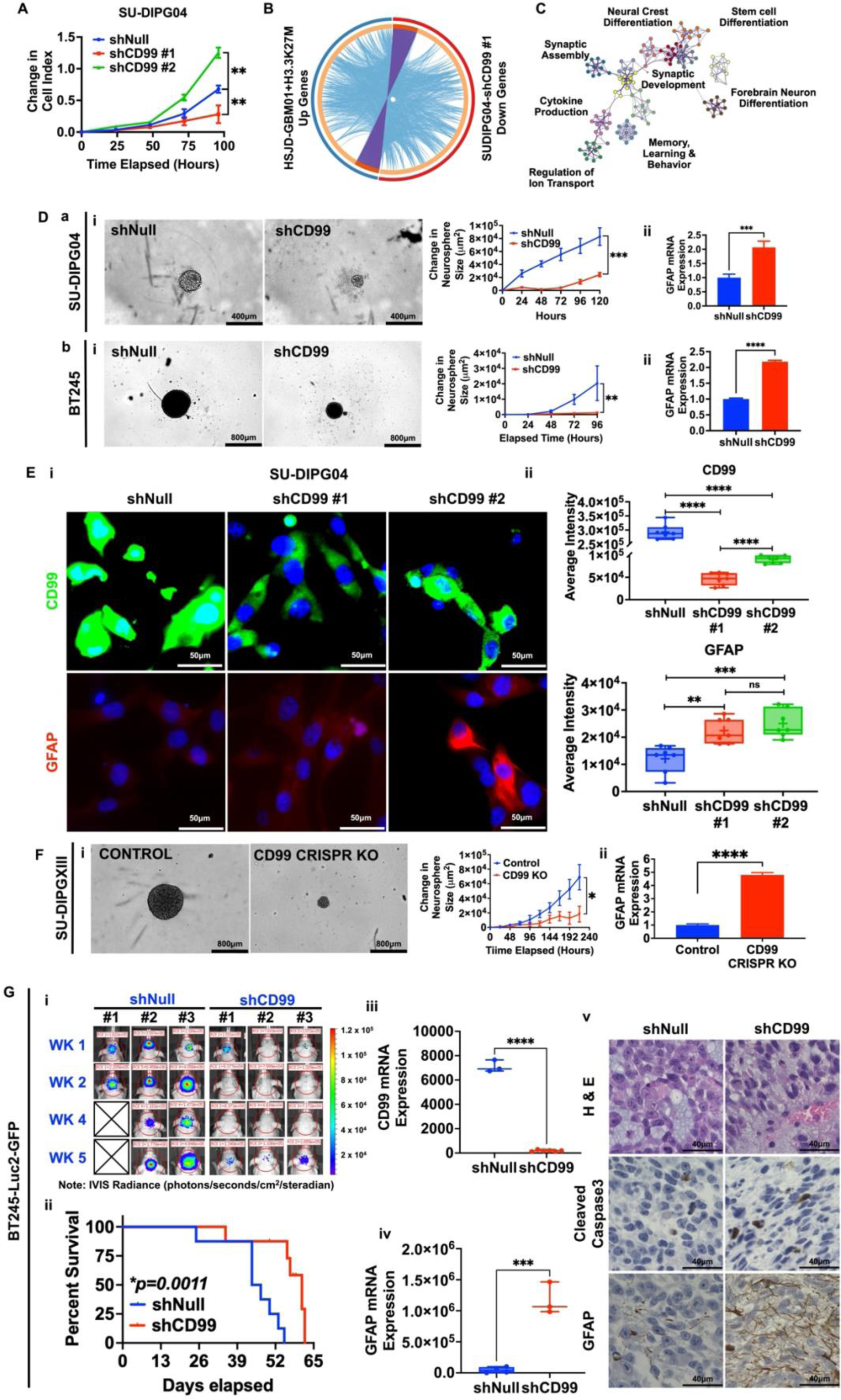
Knockdown or deletion of CD99 restricts the DMG cell growth while inducing tumor cell differentiation. A. shRNA-mediated depletion of CD99 in SU-DIPG04 cells, using different constructs, showed a decrease in their proliferation as measured by xCELLigence. (**p<0.01) B. Gene ontology analysis found that several genes that were downregulated in HSJD-GBM01+K27M were also upregulated in SU-DIPG04 shCD99 #1. C. Metascape analysis of upregulated genes in SU-DIPG04 shCD99 #1 (Figure 2B) shows that these genes are involved in critical neuronal pathways including neuron differentiation, synaptic assembly and development, and cytokine production. D. Neurosphere growth was diminished by shRNA-mediated knockdown of CD99 (a) SU-DIPG04 and (b) BT245. (i) Representative images of neurosphere from incucyte. The adjacent graph shows the quantification of neurosphere size over time. (ii) shRNA-mediated knockdown of CD99 in DMG cells increased the mRNA expression of the differentiation marker, GFAP. (**p<0.01, ***p<0.001) E. (i) Representative IF images show that shRNA-mediated knockdown of CD99 in SU-DIPG04 enhances their differentiation as measured by an increase in GFAP. The nuclei are stained with DAPI (blue), CD99 (green), and GFAP (red). (ii) The quantification of CD99 and GFAP from at least five high-power fields of images. F. Crispr-CAS9 deletion of CD99 in SU-DIPGXIII cells slowed neurosphere growth and increased differentiation. (i) Representative neurosphere images from incucyte. The adjacent graph shows the quantification of neurosphere size over time. (ii) Increase in the mRNA expression of the differentiation marker, GFAP. (*p<0.05) G. shRNA-mediated depletion of CD99 in BT245-Luc2-GFP delayed the tumor initiation and establishment *in vivo*. (i) Representative *in vivo* bioluminescence images (BLI) of the Luc2-expressing DMG tumor cells after injection of the DMG tumor cells in the pons. The scale bar adjacent to the image displays bioluminescence counts (photons/second/cm^2^/steradian). (ii) Kaplan-Meier survival analysis of mice engrafted with CD99-sufficient (shNull, n = 8) and CD99-deficient (shCD99, n = 8) DMG cells. Log-rank (Mantel–Cox) test was used to calculate statistical significance (*p=0.0011). Expression of CD99 mRNA (iii) and GFAP mRNA (iv) in mice tumors from (ii) collected at end-point by RT-qPCR. (v) Representative IHC slides staining H&E and cleaved caspase3 and GFAP in tumors collected at the endpoint. Data shown as the mean ± SEM. (See also **Supplementary Figure S3 and Tables S4 and S5**)

### CD99 knockdown decreased DMG cell self-renewal, induced cell differentiation, and impaired tumor growth

Studies have shown that CD99 is a tumor-associated antigen overexpressed in AML tumor cells, particularly in the leukemic stem cell population (23,29–31). In EWS, with EWS-FLI fusion, previous reports demonstrated that CD99 is required for EWS tumorigenesis, which blocks neural differentiation of EWS tumor cells and observed a modest overlap in the EWS-FLI and CD99 gene regulatory signatures (32). To better understand the potential overlap between gene regulation by the oncohistone K27M mutation and CD99 in DMGs, RNA sequencing (RNA-seq) was conducted on paired samples from SU-DIPG04-shNull and SU-DIPG04-shCD99 DMG cells (**Supplementary Table S5**). Differential gene expression analysis was then performed on the resulting data, which were compared to that of HSJD-GBM01 and HSJD-GBM01+K27M paired analysis (Figure 1A). This analysis allowed us to identify that many genes suppressed by the K27M mutation in HSJD-GBM01 cells were re-activated by CD99 depletion in DMG cells (**Figure 2B**, purple band). Subsequent Gene ontology analysis using Cytoscape (https://cytoscape.org) revealed that most of the altered genes were involved in the induction of neurogenesis, proapoptotic networks, and differentiation pathways **(Figure 2C)**. To evaluate the specific effect of CD99 on DMG cell self-renewal and stem cell markers, a neurosphere assay was performed using CD99 knockdown K27M^+^SU-DIPG04 cell lines. Knockdown of CD99 in DMG cells (1) suppressed neurosphere growth, as shown in representative figures of neurospheres with adjacent graphs showing the quantitation of growth index (**Figure 2Da(i), Db(i)),** (2) decreased the expression of stem cell markers, Nestin and Olig2 (**Supplementary Figures S3D**), while simultaneously inducing the expression of the differentiation marker, GFAP, suggesting a shift from a stem/progenitor state to a more differentiated state (**Figures 2Da(ii), Db(ii), 2E**). Consistent with these findings, CD99 inhibition decreased the stem cell phenotype of SU-DIPG04 cells, as measured by a significant decrease in aldehyde dehydrogenase activity (ALDH), a marker of cancer stem cells (**Supplementary Figures S3E**). In parallel, CRISPR-CAS9 mediated deletion of CD99 from DMG cells (**Supplementary Figure S3F, G**) showed a significant decrease in tumor cell proliferation (**Figure 2F(i)**) and a substantial increase in the differentiation marker GFAP (**Figure 2F(ii)**). These results suggest that K27M induces the expression of CD99, which then signals downstream genes to maintain a stem cell phenotype, thereby preventing differentiation in transformed cells.

The *in vitro* effects of CD99 were also evaluated *in vivo.* shCD99-knockdown BT245 K27M**^+^**DMG cells displayed delayed tumor establishment (**Figure 2G(i)**) and a moderate but significant increase in animal survival compared to shNull cells, which showed rapid tumor establishment and growth (**Figure 2G**(ii), Supplementary Figure 3H**(i)**). Tumors were collected at the endpoint from these mice, and tumoral CD99 expression in the tumor homogenates was tested using qRT-PCR. At the mRNA level, CD99 knockdown tumors maintained significantly lower levels of CD99 than control shNull tumors (p<0.05, **Figure 2G(iii)**). We then examined the expression of GFAP and the active apoptotic marker cleaved caspase 3 in these tumor xenografts by IHC. As indicated in **Figure 2G(iv-v)** and **Supplementary Figures 3H(ii-iii)**, there was a significant increase in GFAP and cleaved caspase 3 in shCD99 tumors compared to shNull tumors. These data suggest that the K27M mutant and the CD99 gene signature overlap, thus co-operatively inhibiting stem cell differentiation and promoting tumor growth. As multiple oncogenic signaling pathways are involved in tumor cell proliferation and growth, we first investigated the possible mechanism of the oncogenic role of CD99 in DMG tumors.

### Upregulated CD99 plays an oncogenic role, in part via integrin-linked kinase-1 (ILK1)

Our data and those of others have shown that CD99 plays a crucial role in preventing tumor cell differentiation, specifically in DMGs. However, the mechanism through which CD99 mediates signal transduction remains unclear. Previous studies have demonstrated that CD99 may regulate β-integrin in T-cells and monocytes (33). These results prompted us to evaluate whether CD99 transduces signals via an integrin-mediated pathway. Integrin-linked kinases (ILK) play a key role in cell proliferation, adhesion, differentiation, and angiogenesis (34,35). One of the integrin-linked kinase, ILK isoforms, ILK1, has been previously shown to regulate oncogenic function in various tumor cell lines (36). To understand whether there is an association between CD99 and ILK1, the changes in ILK1 expression were tested in CD99-modulated DMG cells (Figures 1A-C). We found that similar to CD99, the expression of ILK1 increased in the neoplastic cells upon expressing the K27M mutant, while no significant changes were observed in the K27M-modified NHA cells (**Figure 3A-B**). DMG patient tumor tissues and cell lines highly expressed the ILK1 protein (**Figure 3C-D**). To identify the relationship between the expression of CD99 and ILK1, CD99 knockdown DMG cells were probed for ILK1 protein expression. Immunoblot analysis showed that shRNA-mediated knockdown of CD99 decreased ILK1 expression compared to that in shNull-transduced cells (**Figure 3E**). We also found a loss of CD99 in CRISPR-cas9 K27M KO BT245 cells (Figure 1C); therefore, similar immunoblotting analyses were performed using these cells to explore the K27M-CD99-ILK1 regulatory axis in DMGs. As depicted in **Figure 3F**, deletion of K27M in DMG cells resulted in the downregulation of both CD99 and ILK1 protein, suggesting a possible role of the K27M-CD99-ILK1 axis in the DMG tumorigenesis.

**Figure 3:**
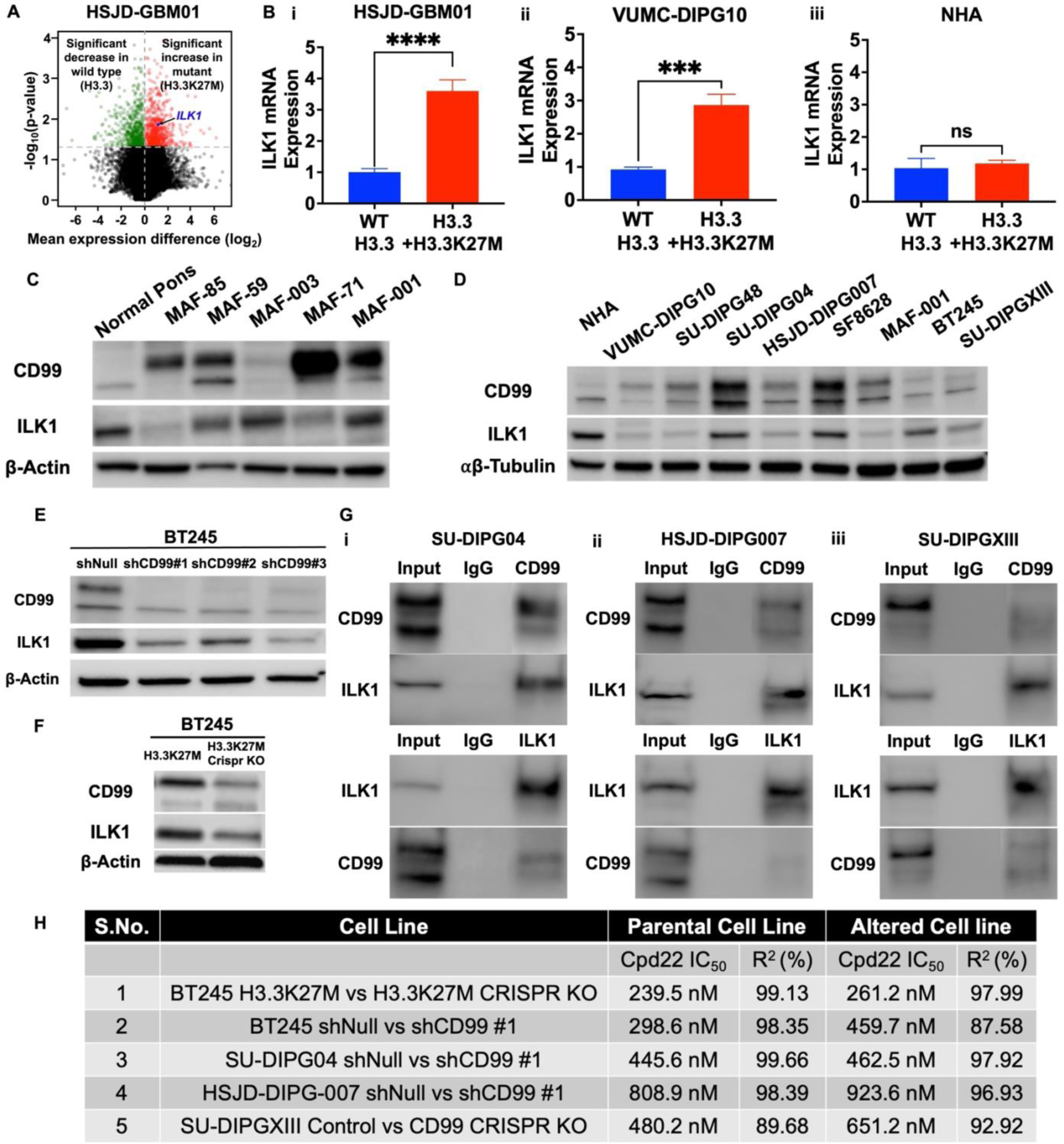
The K27M-CD99-ILK1 is an oncogenic axis in DMG. A. Volcano plot showing the statistical significance as -log_10_(p-value) plotted against the log_2_(fold change) between the genes expressed by HSJD-GBM01 transduced with K27M mutant at the H3.3 locus and the unmodified HSJD-GBM01 (control) cells. HSJD-GBM01 transduced with K27M mutant demonstrated an increase in Integrin-linked kinase 1 (ILK1, indicated with blue arrow) in comparison to the control cells. B. ILK1 mRNA expression in (i) HSJD-GBM01, (ii) VUMC-DIPG10, and (iii) NHA, transduced with H3.3K27M. (ns = not significant, ***p<0.001, ****p<0.0001) C. ILK1 protein, similar to CD99, is highly expressed in DMG patient tumors as compared to normal pons (**C**) and cultured DMG cell lines as compared to normal pons (**D**) by western blotting. E. Western blot showing shRNA-mediated knockdown of CD99 in BT245 cells decreased ILK1. F. Western blot showing Crispr-CAS9-mediated deletion of H3.3K27M in BT245 cells decreased ILK1. G. Coimmunoprecipitation assay showed consistent and modest binding of ILK1 and CD99 proteins in (i) SU-DIPG04, (ii) HSJD-DIPG007, and (iii) SU-DIPGXIII using either pulldown antibodies for CD99 and ILK1. H. The changes in IC_50_ values of the ILK1 inhibitor, cpd22, in CD99-modulated DMG cell lines show that CD99 is an upstream regulator of ILK1 as measured by the MTS assay. Data shown as the mean ± SEM. (See also **Supplementary Figure S4 and Table S4**)

To investigate whether the changes in these two proteins are cooperative, Co-IP experiments were performed to detect whether a direct or distinct interaction between CD99 and ILK1 exists in DMG cells. Co-immunoprecipitation combined with CD99 pull-down and western blot analysis of ILK1 demonstrated the binding of CD99 to ILK1 in DMG cell lines. Similar experiments were performed with pull-down using the ILK1 antibody to validate this interaction further, and pull-down extracts were used to detect CD99 (**Figure 3G**). Similarly, moderate binding of ILK1 to CD99 was noted in the ILK1 pull-down proteins, suggesting protein-protein binding between CD99 and ILK1 (**Figure 3G**). Proteins were not detected in the extracts where the respective IgG controls were used instead of CD99- or ILK1-specific antibodies.

We then examined whether these two proteins functionally interact and whether ILK1 functions as a downstream signal for CD99. A small-molecule ILK1-specific inhibitor, cpd22 (EMD Millipore #407331), has been previously described (37). To test our hypothesis that CD99 exerts its oncogenic function through activating ILK1 in DMGs, we used cpd22 and determined its IC_50_ value in CD99-rich and CD99-modulated DMG cell lines. The IC_50_ value of cpd22 was consistently high in CD99 knockdown and CD99-deleted DMG cells compared to control CD99-rich cells (**Figure 3H and Supplementary Figure S4**). These studies demonstrate the possible role of the oncogenic axis K27M-CD99-ILK1 in DMG tumor cell proliferation.

### A novel anti-CD99 antibody with a membrane-proximal binding epitope showed antitumor efficacy against DMG tumor cells *in vitro*

Because CD99 is a cell surface antigen, we hypothesized that antibodies that block CD99 can inhibit DMG tumor growth. Using informatics tools and existing literature, a new 15 amino acid motif on the extracellular domain of CD99 was designed. A new monoclonal, chimeric CD99 antibody with a membrane-proximal binding epitope was developed (**Figure 4A**). We used this peptide to generate a suite of monoclonal hybridomas and screened them for the optimal clone. We identified one clone, 10D1, with an excellent affinity for binding to cell surface CD99 protein and blocking CD99 expression in DMG cells, as measured by flow cytometry and IHC (data not shown). The 10D1 clone was sequenced and a recombinant antibody was generated on a human IgG4 scaffold (recombinant human 10D1 antibody). The dissociation constant (KD) of 10D1 was measured using the Octet Biacore assay system (38) and found to be in the low nanomolar range in line with other clinically relevant antibodies (**Figure 4B**, **Supplementary Figure S5A**), revealing the high binding affinity of 10D1. This novel monoclonal CD99 antibody, 10D1, showed effective binding to human CD99 expressed on a variety of DMG cell lines as measured by flow cytometry (**Figure 4C**), and its enhanced and specific binding to CD99 on the surface of tumor cells is depicted in **Figure 4D** by IHC. CD99 protein expression detected using 10D1 in the K27M**^+^**DMG tumors mirrored that shown in Figure 1G by western blotting, and 10D1 predominantly detected the long form of CD99 (32kDa) (**Figure 4E, Supplementary Figure S5B**). In addition, as measured by flow cytometry, the ability of the 10D1 antibody to significantly block CD99 in DMG cells is shown in **Figure 4F and Supplementary Figure S5C**.

**Figure 4:**
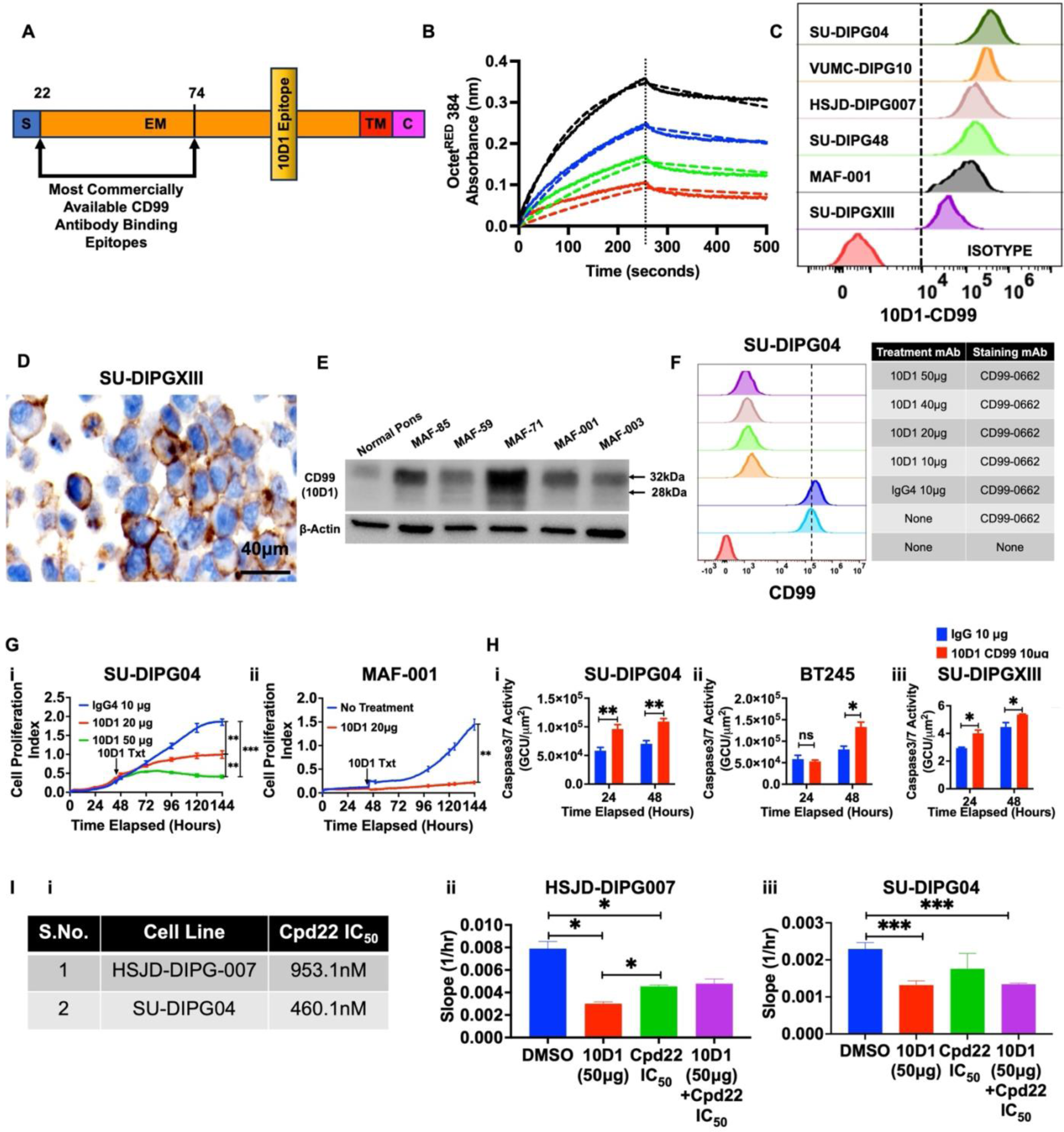
10D1, a novel chimeric monoclonal antibody against CD99, demonstrates high binding affinity to CD99 antigen and inhibits DMG cell growth. A. Schematic showing the human CD99 protein and the approximate location of the 10D1 antibody epitope compared to some of the commercially available CD99 antibody binding epitopes. B. Kinetics of peptide antigen interacting with 10D1 human IgG4/light chain1 using SA biosensor. OctetRED 384 monitored the association and dissociation of this peptide antigen. C. Flow plot showing the detection of CD99 in cultured DMG cells using the 10D1. D. Representative IHC image for staining of CD99 in SU-DIPGXIII cells using the 10D1. E. Detection of CD99 (long isoform) by the 10D1 antibody in DMG patient tumors and normal pons. F. 10D1 significantly blocked CD99 in DMG cells as detected by CD99-0662 antibody even at its low concentrations. G. 10D1 significantly inhibits the proliferation of (i) a DMG tumor line, SU-DIPG04, and (ii) a primary cultured cell line, MAF-001 as measured by xCELLigence. (**p<0.01, ***p<0.001) H. 10D1 significantly induces apoptosis in (i) SU-DIPG04, (ii) BT245, and (iii) SU-DIPGXIII, as measured by the green cleaved caspase3/7 activity on an incucyte. The number of green cells increased significantly 24hrs and 48hrs after treatment. (ns = not significant. *p<0.05, **p<0.01) Data shown as the mean ± SEM. (See also **Supplementary Figure S5**)

We then sought to determine the *in vitro* antitumor efficacy of 10D1 against DMG tumors. Our data showed that 10D1 potently suppressed DMG cell proliferation (**Figure 4G**). A concentration-dependent increase in apoptosis was observed when DMG cells were treated with 10D1 antibody, as measured by an increase in cleaved caspase 3/7 using Incucyte real-time systems at different time points (**Figure 4H**). Conversely, no significant change in cell proliferation of either NHA or induced pluripotent stem cells (iC7-2) expressing low levels of CD99 *in vitro* treated with varying 10D1 concentrations was noted as measured by XCELLigence (**Supplementary Figure S5D-F)**. *These results imply that 10D1 specifically kills high CD99-expressing tumor cells and causes limited toxicity in low CD99-expressing normal cells*.

We then investigated whether ILK1 function downstream of the CD99 pathway could be blocked using 10D1. To test this, we performed rescue experiments; DMG cells were treated with the 10D1 antibody (50μg), cpd22 (IC_50_) alone, or 10D1 followed by cpd22 (IC_50_). We examined the changes in cell proliferation rate (slope) using XCELLigence using two DMG cell lines. While both 10D1 or cpd22 treatment alone inhibited tumor cell proliferation compared to control-treated (DMSO) cells, blocking CD99 first with 10D1 rescued cells from subsequent cpd22-induced cell death (**Figure 4I, Supplementary Figure S5G**). As the specificity of cpd22 in inhibiting ILK1 has been well documented (37), these findings suggest that ILK1 acts as a downstream signaling molecule that plays a functional role in CD99-driven oncogenic events in DMG. While this is an exciting finding, it warrants further detailed *in vivo* studies using the combination of 10D1 and cpd22 effects on DMG tumor growth.

### 10D1 can cross the blood-brain barrier (BBB) and exert an anti-tumor effect against DMGs *in vivo*

Antibodies are potent anticancer therapeutic tools (39). However, targeting brain tumors is particularly challenging because of the presence of the BBB. To test whether CD99 antibodies can cross the BBB to produce a therapeutic effect, *in vivo,* experiments were performed following the treatment protocol shown in **Figure 5A**. Luciferase-expressing BT245 cells were established in NSG mice, as confirmed by IVIS imaging. Animals were randomized and treated intravenously (IV) with IgG4 or 10D1 (n=5/group) at (8mg/kg/day). A dose of 8 mg/kg was effective and safe in dose-dependent studies (data not shown). Three hours after the infusion of the 3^rd^ final dose of the antibodies (IgG4 or 10D1), perfused brain tissues were collected and analyzed for the presence of IgG4. Significant binding of IgG4 was observed in 10D1 antibody-treated tumors, but not in IgG4-treated tumors (**Figure 5B**). Next, to test the anti-tumor efficacy of 10D1, following the protocol in **Figure 5C**, animal cohorts (n=3) that received a total of 6 doses of either IgG4 or 10D1 IV at 8 mg/kg were monitored for changes in the tumor volume by IVIS imaging and by magnetic resonance imaging (MRI). CD99 antibody-treated animals showed a significant decrease in tumor burden in the pons compared to that in IgG control-treated mice (**Figures 5D-E**). IHC analysis of the brain tissue sections collected at this time point showed tumor regression (**Figure 5F)**, as evidenced by reduced K27M and CD99 staining in 10D1 treated tumors. *These data suggest that the 10D1 anti-CD99 antibody can cross the BBB and exert its antitumor effect on DMG tumor growth*.

**Figure 5:**
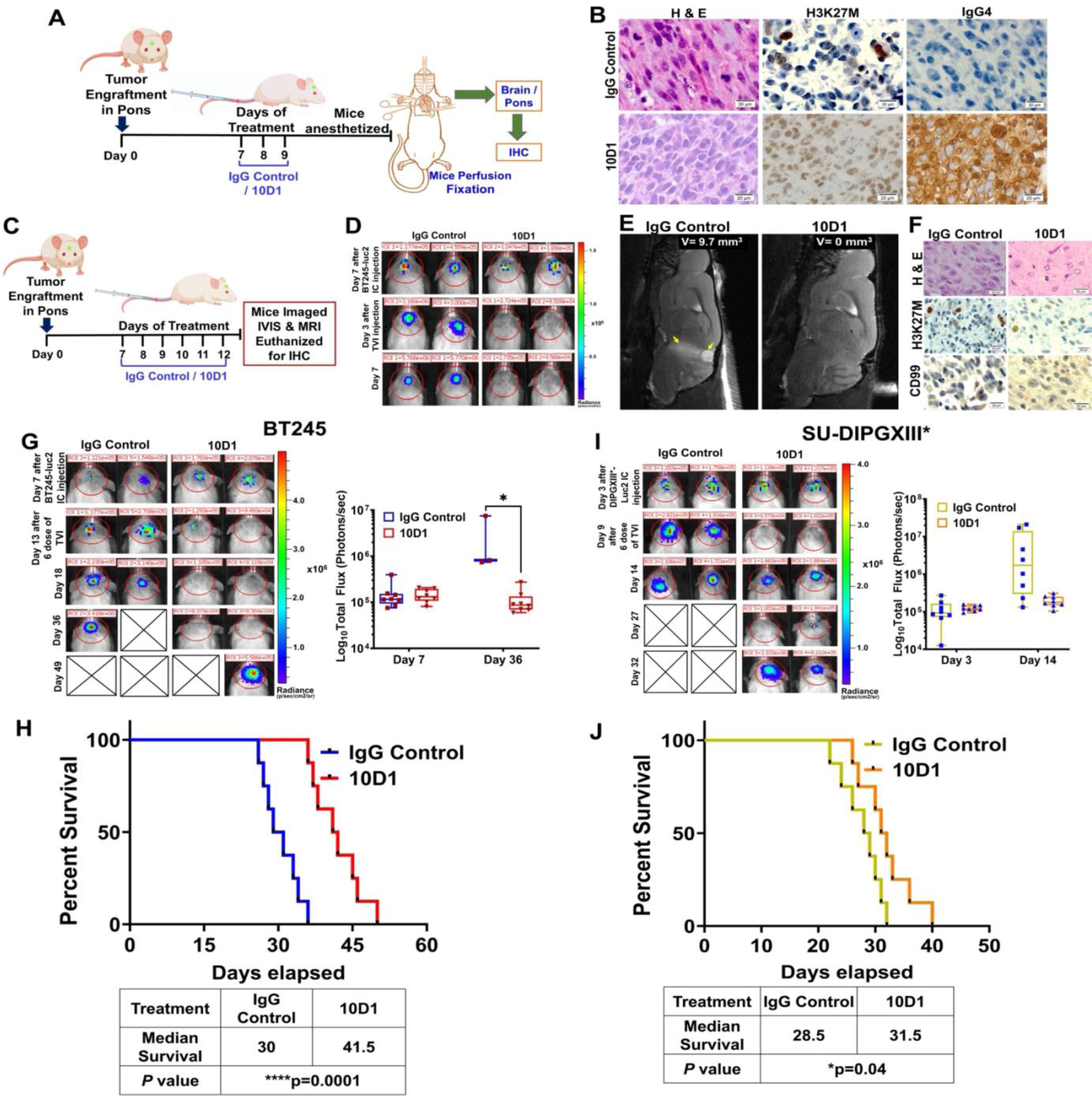
Intravenous (IV) delivery of 10D1 antibody inhibits the progression of DMG tumors *in vivo*. *A-B : Pilot experiments to find if 10D1 could cross the blood-brain barrier (BBB)* A. Schematic diagram of a pilot study to investigate the brain penetrance of 10D1 antibody in BT245 pons tumor-bearing mice. The mice were infused with 8mg/kg of either IgG4 or 10D1 antibody for 3 consecutive days. B. IHC staining images of H&E, H3K27M, and IgG4 in the perfused brain tissues collected 24 hrs after the final infusion of respective antibodies, as shown in A. The positive IgG4 staining in the tumor tissues exposed to 10D1 suggested the ability of 10D1 to cross BBB. *C-F: Experiment to evaluate the antitumor efficacy of infused 10D1 against DMG*. C. Schematic showing the protocol followed to evaluate the anti-tumor efficacy of 10D1 protein using BT245 tumor-bearing mice. The mice were infused with 8mg/kg of either IgG4 or 10D1 antibody for 6 consecutive days. D. Representative BLI showing the changes in the tumor burden in pons tumor-bearing mice treated with the respective antibody treatments as given in C. E. Representative MRI images of the mouse brain taken 24 hrs after 6 doses of the treatment showed complete clearance of the tumor in the 10D1-infused mouse while the tumor continued to grow in the IgG4 control mouse (yellow arrows). F. After the MRI images were done as in E, the brain tissues were collected and analyzed for the presence of a tumor as envisioned by H&E and H3K27M staining. There is a decrease in the size of residual tumor after 10D1 treatment suggesting the antibody could cross the BBB and demonstrate antitumor efficacy. *G-H: Experiments to evaluate the effect of 10D1 on animal survival using the BT245-Luc2 model*. G. Experiments using a large cohort of animals were performed. Representative BLI of pons BT245 tumor-bearing mice before and after treatment with the IgG4 control and 10D1 antibody following the protocol given in supplementary figures 6A. The adjacent plot shows changes in flux intensity before and after the respective treatments on the specified days. (*p<0.05) H. Kaplan Meier survival curve of pons BT245 tumor-bearing mice shows that 10D1 treatment increased the survival compared to control-treated mice. Log-rank (Mantel–Cox) test was used to compare groups (****p<0.0001). n = 7 to 9 mice per group. *I-J: Experiments to evaluate the effect of 10D1 on animal survival using the SU-DIPGXIII*-Luc2 model*. I. Representative BLI of pons SU-DIPGXIII* tumor-bearing mice before and after treatment with the IgG4 control and 10D1 antibody following the protocol given in supplementary figures 6B. The adjacent plot shows changes in flux intensity before and after the respective treatments on the specified days. (*p<0.05). J. Kaplan Meier survival curve of pons SU-DIPGXIII* tumor-bearing mice showed that 10D1 treatment modestly increased the survival compared to control-treated mice. Log-rank (Mantel– Cox) test was used to compare groups (*p<0.05). n = 7 to 9 mice per group. The respective animal median survival is shown in the table below. Data represented as the mean ± SEM. (See also **Supplementary Figure S6**).

Following these pilot studies, similar *in vivo* experiments were performed using a larger cohort of animals established with two different DMG tumor models, BT245-Luc2 and SU-DIPGXIII*-Luc2 tumors, as shown in **Figures 5G-J** and **Supplementary Figures S6A-J**. In the BT245 mouse model, treatment with 10D1 significantly reduced the tumor burden and significantly increased animal survival (**Figure 5G-H, Supplementary Figure S6B-C**). In addition, mice treated with 10D1 gained weight compared to the IgG control, suggesting improved overall health (**Supplementary Figure S6D**). At the animal study endpoint, tumoral analysis of CD99-treated tumors in the BT245-Luc2 model demonstrated an increase in cleaved caspase 3 compared to controls, further suggesting a therapeutic effect (**Supplementary Figure 6E**). In the SU-DIPGXIII*-Luc2 model, following the treatment protocol in **Supplementary Figure 6F**, although there was an initial reduction in the tumor burden with the infusion of 10D1, the tumors relapsed quickly, leading only to a modest but significantly increased animal survival (**Figures 5I-J, Supplementary Figures 6G-H**). In addition, animal body weight increased during the 10D1 treatment time frame (**Supplementary Figure 6I**). The failure to observe enhanced antitumor efficacy and animal survival with 10D1 treatment in the SU-DIPGXIII*-Luc2 model could be due to the aggressive growth properties of this tumor type (27). Irrespective, at the animal study endpoint, tumoral analysis of CD99-treated tumors in the aggressive SU-DIPGXIII*-Luc2 model demonstrated a similar increase in cleaved caspase 3 compared to controls, further establishing a therapeutic effect (**Supplementary Figure 6J**). These data confirmed that the systemic administration of 10D1 could cross the BBB and inhibit tumors in orthotopic pontine DMG xenografts.

### Locoregional (intrathecal) delivery of 10D1 enhances animal survival

As an alternative delivery route for 10D1, we questioned whether intrathecal (IT) delivery could further enhance the antitumor efficacy of 10D1 *in vivo* animal DMG models. To test this, a single low dose of 10D1 or IgG4 antibody at 1 mg/kg was delivered locoregionally to the 4^th^ ventricle area in established pontine-tumor-bearing NSG mice following the protocol shown in **Figure 6A** (BT245-Luc2). Locoregional administration of 10D1 significantly attenuated DMG tumor growth (**Figure 6B, Supplementary Figures S7A-B**) On day 7, antibody-treated mice showed a drastic decrease in tumor burden after 10D1 treatment and increased body weight (**Supplementary Figure S7C**). In contrast, the tumor continued to grow in control-treated mice, with a decrease in animal body weight. Kaplan-Meier analysis comparing IgG control and 10D1 antibody delivered at a low dose (1 mg/kg) to the CNS showed that 10D1 treatment significantly enhanced the survival of BT245-Luc2 mice (**p=0.0018, **Figure 6C**).

**Figure 6:**
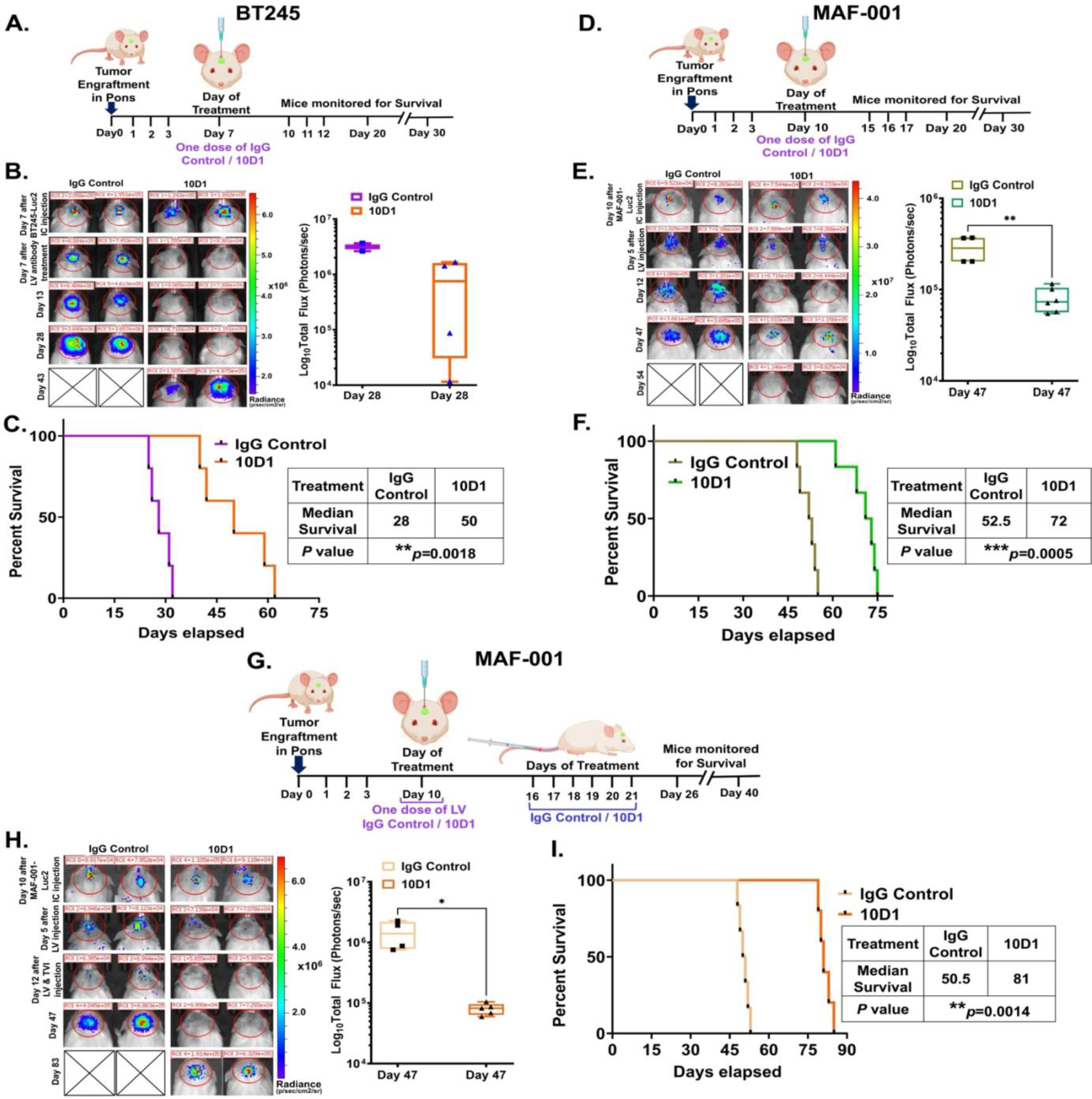
Intrathecal (IT) delivery of a single low dose of 10D1 prolonged DMG xenograft survival. *A-C : Effect of IT delivery of a single dose of 10D1 in BT245-Luc2 model*. A. Schematic for treatment protocol followed for BT245 model. B. Representative BLI of mice before and after specified treatments in the BT245 xenograft models. The adjacent plots show flux values before and after treatments on the specified days. C. Kaplan Meier survival curve shows that 10D1 treatment increased the survival of pons BT245 tumor-bearing mice compared to control-treated mice. Log-rank (Mantel–Cox) test was used to compare groups (**p<0.01) n = 5 to 6 mice per group. The animal median survival is shown in the table below. *D-F: Effect of IT delivery of a single dose of 10D1 in a primary cell line, MAF-001-Luc2, model*. D. Schematic for treatment protocol followed for MAF-001 PDX model. E. Representative BLI of mice before and after specified treatments in the MAF-001 PDX model. The adjacent plots show flux values before and after treatments on the specified days (**p<0.01). F. Kaplan Meier survival curve shows that 10D1 treatment increased the survival of pons MAF-001 tumor-bearing mice compared to control-treated mice. Log-rank (Mantel–Cox) test was used to compare groups (***p<0.0005). n = 5 to 6 mice per group. The animal median survival is shown in the table below. (See also **Supplementary Figure S7**). *G-I: Effect of combined delivery modes: IT delivery of a single dose of 10D1 followed by intravenous infusion of 10D1 for 6 consecutive days in the MAF-001-Luc2 Model*. G. Schematic for treatment protocol followed for MAF-001 PDX model. H. Representative BLI of mice before and after specified treatments in the MAF-001 PDX model. The adjacent plots show flux values before and after treatments on the specified days (*p<0.05). I. Kaplan Meier survival curve shows that 10D1 treatment increased the survival of pons MAF-001 tumor-bearing mice compared to control-treated mice. Log-rank (Mantel–Cox) test was used to compare groups (**p<0.01). n = 5 mice per group. The animal median survival is shown in the table below. Data represented as the mean ± SEM. (See also **Supplementary Figure S8**).

Next, we tested the anti-tumor efficacy of IT delivery of 10D1 in our newly established patient-derived orthotopic pons tumor model, MAF-001-Luc2, using the protocol shown in **Figure 6D**. Administration of 10D1 decreased the tumor burden of the patient-derived xenografts, a similar effect as seen with the cell line xenografts (**Figure 6E, Supplementary Figures S7D-E**). MRI performed six days after a single dose of antibody treatment further confirmed the decrease in tumor burden with 10D1 treatment (**Supplementary Figure S7F).** Kaplan-Meier survival analysis showed that 10D1 treatment in the primary tumor model also significantly enhanced the survival of MAF-001 xenograft-bearing mice (\*\*\**p*=0.0005, **Figure 6F**). Complete Blood Count (CBC) analysis of blood samples collected from a cohort of these mice at the study endpoint showed no evidence of hematological toxicity due to either antibody treatment (**Supplementary Figure S7G**).

Based on the results of the 10D1 treatment, we sought to determine whether the anti-tumor latency of 10D1 could be increased by combining both routes of administration: IT followed by IV. To test this, pons tumor-bearing NSG mice established with patient-derived primary MAF-001-Luc2 DMG tumor cells were treated using the method shown in **Figure 6G**. In this method, a single dose of IgG Control or 10D1 antibody (1 mg/kg) was delivered via the 4^th^ ventricle, followed by IV injection (8 mg/kg) for six consecutive days of treatment. The BLI images and quantitation of the tumor burden from these images revealed that the sequential delivery of 10D1 increased the anti-tumor latency compared to either treatment alone (**Figure 6H and Supplementary Figures S8A-B**), with a significant increase in animal body weight (**Supplementary Figure S8C**). IT followed by IV infusion of 10D1 enhanced animal survival by an additional 12.5% compared to IT treatment alone in the MAF-001-Luc2 animal model (**Figure 6I**), while the hematological parameters analyzed showed no significant changes in either treated group of mice due to the administration of antibodies consecutively through two delivery routes (**Supplementary Figure S8D**).

### 10D1 combined with radiation treatment (RT) exerts potent anti-tumor efficacy in DMG

The current standard of care for patients with DMG is RT, which is only palliative. Therefore, improving the efficacy of RT is likely to significantly affect the outcomes of patients with DMG. We hypothesized that combining 10D1 with radiation would improve the antitumor efficacy of radiation. Hence, we first examined radiation-induced changes in the cell surface expression of CD99 in DMGs using flow cytometry. DMG cells treated with single bolus RT doses showed a minimal change in CD99 expression (**Supplementary Figure 9A**). To simulate clinical focal fractionated RT (FFRT), DMG cells were exposed to different fractionated doses of radiation *in vitro*. Flow cytometry results showed a significant and consistent increase in CD99 with increasing fractionated radiation doses in K27M^+^DMG cells, but no such changes were observed with that of NHA cells (**Figure 7A**, **Supplementary Figure S9B-C**). Furthermore, in biopsy-derived radiation-naïve SF8628 DMG cells, western blotting analysis showed that while fractionated RT increased CD99 protein levels, it predominantly induced the active long form of CD99 and not the short form (**Figure 7B**).

**Figure 7:**
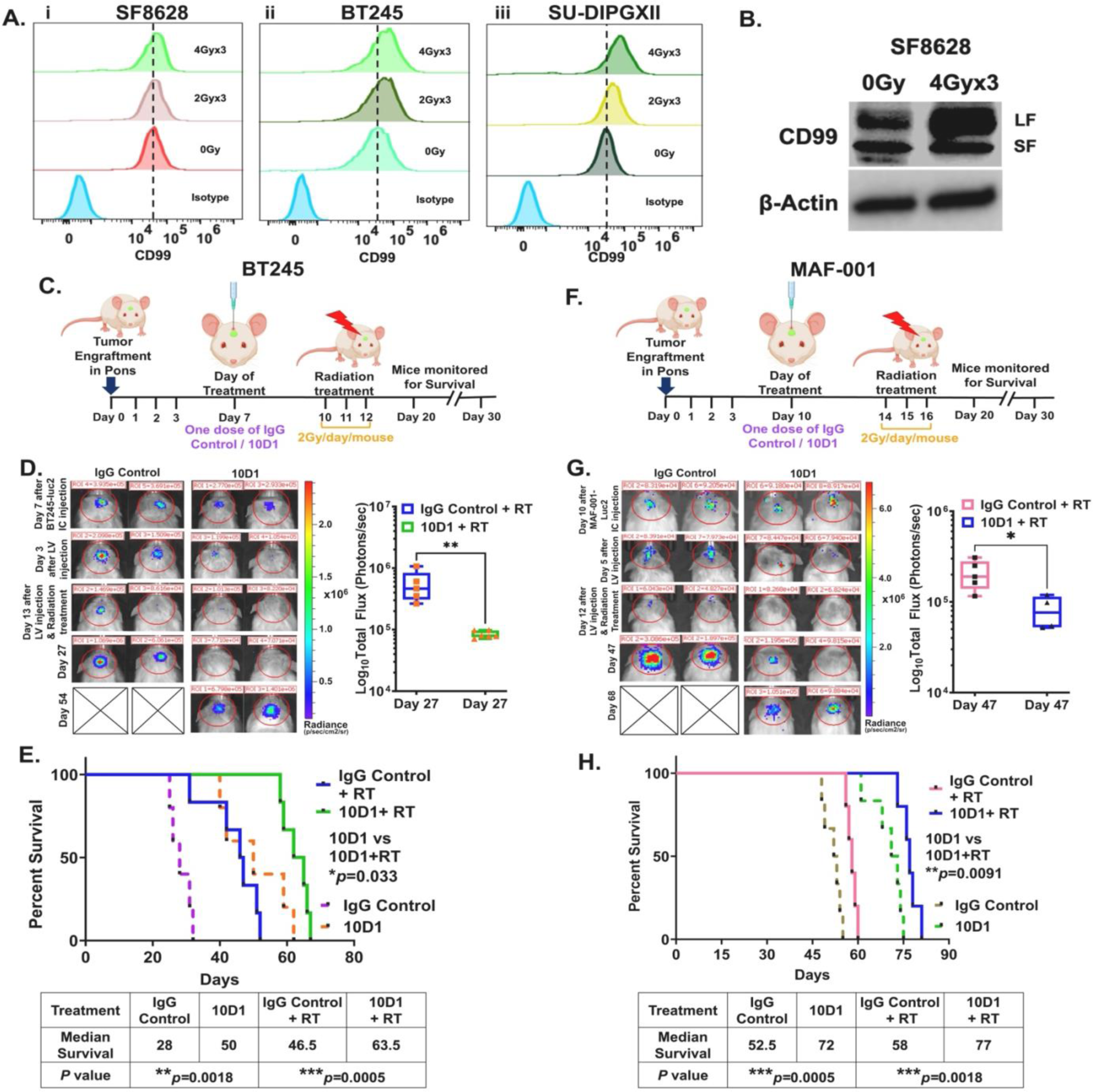
Fractionated radiation increases CD99 in DMG cells and antagonizing CD99 with 10D1 increased the anti-tumor latency of radiation treatment (RT) in DMG xenografts. *A-B: RT increases CD99 in DMG cells in vitro*. A. Representative flow cytometry plots showing the expression of CD99 at 24hrs after they were exposed to RT of 2Gy or 4Gy dose for three consecutive days in (i) SF8628, (ii) BT245, and (iii) SU-DIPGXIII, (quantitation of CD99 expression is given in **Supplementary Figure S9C**). B. A representative western blot shows the increased CD99 protein (only the long form) after SF8628 cells were exposed to RT with a 4Gy dose for three consecutive days. *C-E: IT delivered 10D1, followed by focal fractionated RT (FFRT) increased the anti-tumor latency of RT in the BT245-Luc2 in vivo model*. C. Schematics representing the treatment protocol followed in the BT245 model. D. Representative BLI images of mice after tumor implantation, antibody treatment, and RT. The adjacent plot shows the flux values comparison on day 27 between the control and treatment groups (**p<0.01). E. Kaplan Meier survival curve shows the increase in survival in BT245 tumor-bearing mice treated with a single IT dose of 10D1 followed by RT in comparison to that of control-treated mice. Log-rank (Mantel–Cox) test was used to compare groups (***p<0.0005). n = 5 to 6 mice per group. The median survival of the mice is given in the table below. Dotted lines in the survival curves indicate intrathecal alone treatment. *F-H In vivo effect of intrathecally delivered 10D1, followed by FFRT in MAF-001-Luc2 model*. F. Schematics representing the protocol followed in the MAF-001 PDX model. G. Representative BLI of mice xenografted with luciferase-expressing MAF-001 before and after 10D1 treatment, followed by RT. The adjacent plot shows the flux values comparison on day 47 between the control and treatment groups (**p<0.05). H. Kaplan Meier survival curve shows the increase in survival in MAF-001 tumor-bearing mice treated with a single IT dose of 10D1 followed by RT in comparison to that of control-treated mice. Log-rank (Mantel–Cox) test was used to compare groups (**p=0.0018). n = 5 mice per group. The median survival of the mice is given in the table below. Dotted lines in the survival curves indicate intrathecal alone treatment. Data represented as the mean ± SEM. (Also see **Supplementary Figure S9**)

These results imply that the heightened expression of CD99, especially its active state (long form) after fractionated RT, appears to serve a protective role in DMG tumor cells, shielding them from the detrimental effects of RT. Therefore, we hypothesized that blocking CD99 using 10D1 followed by RT would enhance the anti-tumor efficacy of RT.

To test this hypothesis, we first performed *in vivo* studies by administering 10D1 via IT delivery followed by FFRT in BT245-Luc2 tumor-established pons xenografts (**Figure 7C**). Blocking CD99 with a single dose of 10D1 delivered intrathecally followed by FFRT prolonged the antitumor efficacy of RT alone, as demonstrated by the persistent decrease in tumor burden well beyond the time frame of IgG+FFRT treatment cohorts (**Figure 7D** and **Supplementary Figure S9D-E**) and prolonged animal survival (**Figure 7E**). The decrease in tumor burden with the combination treatment was reflected in the concomitant increase in animal weight (**Supplementary Figure S9F**), and animals succumbed to tumor burden when the tumors relapsed much later after 10D1 treatment was stopped. The enhanced efficacy of the combinatorial treatment was then tested in primary MAF-001 orthotopic xenografts following the protocol shown in **Figure 7F**. Similarly, we observed an increased latency in tumor growth with reduced tumor burden and a significant improvement in animal survival in MAF-001 orthotopic xenografts treated with the combination of 10D1 and FFRT compared to FFRT alone (**Figures 7G-H**; **Supplementary Figures S9G-H**). The CBC analysis of the blood collected at the animal endpoint showed no significant toxicities with combined treatment (**Supplementary Figure S9I**).

Although it is apparent that a single low dose of 10D1 delivered directly to the intrathecal space showed enhanced antitumor efficacy compared to the IV infusion of 10D1 in pons tumor-bearing mice, we tested whether the systemic infusion of 10D1 by IT could also potentially affect the efficacy of FFRT in the aggressive SU-DIPGXIII*-Luc2 xenograft model following the protocol and dosing shown in **Supplementary Figure S10A**. IT delivery of 10D1 followed by FFRT significantly improved the tumor-killing effect of FFRT as evidenced by the persistent reduction in tumor burden and increased animal survival compared to either treatment alone (**Supplementary Figures S10B-D**). Furthermore, we tested whether 10D1 infusion by TV could also potentially alter the efficacy of FFRT in the BT245-Luc2 xenografts following the protocol and the treatment dose shown in **Supplementary Figure S10E**. Systemic delivery alone of 10D1 followed by FFRT also enhanced the tumor cell killing effect of RT alone with long-term persistence of animals with reduced tumor burden and increased survival compared to either treatment alone (**Supplementary Figures S10F-H**). Taken together, these studies, using different DMG pons tumor xenograft studies, the two routes of administration of 10D1 and its combinatorial treatment with FFRT, increasing anti-tumor effect with enhanced survival were seen in the following order of treatments: **IV < IV+FFRT= IT < IV+IT< IT+FFRT (Supplementary Table S6)**.

## Discussion

Although the identification of new agents for DMG therapeutics has long been appreciated, progress has been limited (3,40). This is due to factors such as tumor cell resistance to chemotherapies and failure of effective drugs to cross the BBB and reach the tumor site, while radiation therapy was only palliative as tumors relapsed and grew rapidly after radiation therapy. With the presence of these roadblocks in the effective treatment of DMG tumors, we investigated novel ways to target DMGs. We identified that the oncogenic driver K27M regulates CD99 to partly exert its oncogenic function and since targeting the K27M directly was not effective, targeting CD99 as a new therapeutic pathway to treat K27M^+^DMGS. Warranting further in detail *in vitro* and *in vivo* research studies, the therapeutic applicability of targeting CD99 can be extended to H3WT expressing DMG tumors, as these tumors also express CD99, although to a lesser extent than K27M^+^DMGs.

CD99 is expressed in a variety of normal and tumor cells including Ewing sarcoma (EWS) and Acute Myeloid Leukemia (AML) tumors. CD99 expression is frequently higher in AML stem cells than in their normal hematopoietic counterparts (22,31,41) and is blocked by a monoclonal antibody (mAb) against CD99-induced cytotoxicity in AML stem cells *in vitro* (*22*). Furthermore, anti-CD99 mAbs exhibited antileukemic activity in AML xenografts *in vivo*, demonstrating the potential of anti-CD99 mAbs as therapeutic agents (42). Similarly, Scotlandi et al. extensively investigated the high expression of CD99 and the biological role of CD99 in EWS. Subsequent studies by this group, using an antibody targeting CD99 (0662, mouse Mab), demonstrated that engagement of CD99 significantly inhibits the *in vitro* and *in vivo* growth of EWS tumor cells (43,44). Notably, CD99 engagement did not induce apoptosis in normal mesenchymal cells but potently induced cell death when normal cells were transduced with the EWS-FLI transgene found in EWS (43).

In the past two decades, there has been a significant surge in the development of antibodies for the treatment of human diseases. In recent years, approximately ten antibodies have been approved by the FDA annually ((45) and www.antibodysociety.org/resources/approved-antibodies). Therapeutic antibodies have been successfully used to treat infections, cancers, and other autoimmune diseases (46,47). Because monoclonal antibodies have high specificity, they hold great promise for the treatment of CNS diseases. Owing to its enhanced expression on the DMG cell surface, CD99 is a compelling immunotherapy target. Several monoclonal antibodies that bind to CD99 are available (48,49). Many of these previously characterized anti-CD99 monoclonal antibodies, such as DN16 and 12E7, have binding epitopes that are distally located from the cell surface membrane, except for the 0662 antibody, which has distinct binding epitopes from the other antibodies (33). To our knowledge, there are currently no clinically accessible CD99 antibodies. To address this, we developed a therapeutic CD99 antibody. As previously shown, the efficacy of a therapeutic antibody is largely based on the binding epitope, and epitopes binding closer to the membrane enhancing antibody-dependent cytotoxicity in cells (50), we developed a novel monoclonal therapeutic anti-CD99 chimeric antibody (10D1) with a membrane-proximal epitope that was engineered with a human IgG4 scaffold.

Previous research has shown that one of the major obstacles in treating CNS diseases with antibody-based therapeutics is the presence of the blood-brain barrier (BBB), which restricts the access of the brain to antibodies. The BBB has low permeability, which helps to maintain brain homeostasis but may also prevent therapeutic molecules from entering the brain (51). In brain tumors, the BBB is abnormal, with morphological changes in the barrier and increased barrier permeability due to disrupted endothelial cell junctions (52). Despite these drawbacks, the 10D1 chimeric antibody, when delivered intravenously, showed an increased ability to cross the BBB, reach the pons tumor site, and mediate its therapeutic effects in murine models of DMGs. Similar, but limited, success in using other antibodies has been demonstrated in murine models of brain tumors, including an anti-EGFRvIII antibody in glioblastoma and an anti-CD47 antibody in pediatric brain tumors (53,54). Simultaneously, multiple strategies have been explored to increase antibody delivery to the brain and to treat CNS tumors, including the development of BBB-penetrant bi-specific antibodies. Examples include the development of a bispecific anti-Ang-2/TSPO antibody used in a glioblastoma mouse model showing brain penetrance (55).

On the other hand, recent progress in administering therapeutics directly to the CNS to treat patients with brain tumors using an intraventricularly implanted Ommaya reservoir offers multiple advantages beyond overcoming the BBB. This approach enables repeated administration of small doses of drugs while increasing potency and enhancing safety by reducing systemic toxicity. Our data demonstrated that delivering 10D1 via this clinically relevant mode of direct delivery would further enhance the therapeutic effect of 10D1 against pontine tumors.

Although palliative radiation therapy is the only standard of care, DMGs are resistant to chemotherapy and small-molecule drug inhibitors (3,56). Radiation therapy is an essential modality for many types of cancers. It leads to a plethora of changes in the tumor and tumor microenvironment, resulting in many undesired off-tumor effects. Based on our data, we postulate that, in reaction to focal fractionated radiation therapy (FFRT), DMG cells trigger the expression of CD99 on their cell surface. Although the exact mechanism of this upregulation of CD99 in the DMG after radiation therapy is unknown and beyond the scope of this study, the consequences of elevated CD99 expression have been shown to induce adaptability in adult glioblastoma (GBM) tumors. This adaptability is implicated in tumor recurrence and therapy resistance in these tumors (57). Our study, in conjunction with other studies, indicates that CD99 plays a crucial role in safeguarding tumor cells from radiation-induced harm. By blocking CD99 with 10D1 and administering FFRT, we were able to enhance the *in vivo* antitumor effectiveness of radiation therapy and extend the survival of DMG xenografts. This suggests that this combination therapy has great potential for improving treatment outcomes in patients with DMG. *In vivo* studies show that administering 10D1 via IT or IV routes, in combination with FFRT, enhanced antitumor efficacy compared to individual treatments. Notably, a single low-dose IT injection of 10D1, followed by FFRT, was found to be the most effective method for extending xenograft survival. Together, these preclinical results may guide future clinical applications of IT delivery of 10D1 followed by FFRT as the most effective and safe therapeutic approach for DMG.

While CD99 is implicated in other cancer types and the use of monoclonal antibody targeting CD99 has demonstrated anti-tumor efficacy *in vitro* and *in vivo* models of Ewing sarcoma, GBM, and AML, these findings, to our knowledge, have not yet resulted in clinical trials (32,58–60). However, outcomes from previous studies investigating the oncogenic role of CD99 across diverse tumor models collectively indicate the potential clinical utility of the therapeutic antibody 10D1 in treating conditions beyond DMGs.

## Materials and Methods

### Patient Tumors and Cell lines

Tumor and normal tissues were collected at the Children’s Hospital of Colorado (CHCO) after informed consent was obtained under the approved IRB protocol (COMIRB 95-500). The clinical and molecular characteristics of the cells established from primary tumors, a complete list of cell lines used in this study, cell culture information, and their sources are given in **Supplementary Table S1**. DNA fingerprinting was performed to authenticate the cell lines used, and the fingerprinting profiles are listed in **Supplementary Table S1**.

### Histone and Gene modification in cells

1. *<u>H3.3K27M mutant overexpression</u>*: H3.3 WT cell lines (VUMC-DIPG10 and HSJD-GBM01) and normal human astrocytes (NHA) were transduced with a lentiviral K27M mutant transgene to stably express the K27M mutant transgene or an empty vector. Briefly, approximately 250,000 cells/well were seeded in a 6-well plate. The cells were then transduced with the K27M mutant transgene using 8ng/ml polybrene. These plasmids were a kind gift from Dr. C. David Allis, Rockefeller University, NY (10). Then, the pure population of transduced cells was selected using puromycin. Overexpression of these histone transgenes was confirmed by western blotting of histone proteins collected using acid extraction (61). These modified cell lines have been named the following in this study: HSJD-GBM01+K27M, VUMC-DIPG10+K27M, and NHA+K27M.
2. *<u>H3.3 WT overexpression</u>*: The H3.3K27M mutant cell line, HSJD-DIPG007, was transduced with the lentiviral H3.3 WT transgene or empty vector to stably express the H3.3 wild-type transgene and overexpression of these histone transgenes was confirmed by western blotting of histone proteins This cell line was indicated in this study as HSJD-DIPG007+H3.3WT.
3. *<u>CD99 knockdown (shRNA-mediated)</u>*: The CD99 gene in DMG lines, SU-DIPG04 and BT245, was knocked down using lentivirus derived from short hairpin RNAs (shRNAs) targeting the CD99 gene or non-targeting controls. These plasmids were purchased from the Functional Genomics Core at the University of Colorado Anschutz Medical Campus (shCD99 #1: *TRCN0000057503*; shCD99 #2: *TRCN00000291644;* shCD99 #3: *TRCN00000291708,* shNull: *TRCN SHC202)* Briefly, 1 × 10^6^ cells/well were seeded in a 100 mm plate and transduced with lentiviral particles using 8ng/ml polybrene. Then, the pure population of the transduced cells was selected with puromycin. Knockdown of CD99 was confirmed by western blotting of proteins collected using RIPA buffer extraction. The following model cell lines were used: SU-DIPG04 shCD99 #1, SU-DIPG04 shCD99 #2, SU-DIPG04 shCD99 #3, BT245 shCD99 #1, BT245 shCD99 #2, and BT245 shCD99 #3. The non-targeting controls were named SU-DIPG04 shNull and BT245 shNull.
4. *<u>Generation of CD99 KO DMG cells (CRISPR/CAS9-mediated)</u>*: CD99 was deleted using the CRISPR-CAS9 knockout protocol, as previously described (62). Briefly, single guide RNAs (sgRNAs) targeting the CD99 gene were obtained from Integrated DNA Technologies (IDT, Hs.Cas9.CD99.1.AB). CRISPR-CAS9 knockout in SU-DIPGXIII cells was performed using the Amaxa P3 Primary Cell 4D-Nucleofector X Kit L (Lonza, V4XP-3012). First, the ribonucleoprotein (RNP) mixture was prepared by mixing the sgRNA (100μM) with the Alt-R Cas9 enzyme (62μM) in 1X PBS. After washing SU-DIPGXIII (∼10×10^6^) cells in PBS, the cells were resuspended in 240μl P3 Primary Cell Solution (Lonza). 15μl Alt-R Cas9 enhancer (IDT, #1075915) and 90μl RNP mixture were added to the cell suspension. After transferring the cell suspension into the nucleofection cuvette, nucleofection was performed on a 4D-NucleofectorTM X unit. The CD99 negative cells were sorted using an XP70 cell sorter, in the Flow Cytometry Core at the University of Colorado Anschutz Medical Campus. Then, a single cell was plated in each well of a low-attachment 96 well plate to identify specific clones with the deletion of CD99. The CD99-negative clones were used for further studies. The deletion of CD99 in the selected clone was evaluated by RT-qPCR using Qiazol ® extraction and is represented as SU-DIPGXIII CD99 KO. The non-targeting control was named SU-DIPGXIII Scramble.
5. <u>K27M mutant KO DMG cells (CRISPR/CAS9-mediated)</u>: Dr. Nada Jabado of McGill University, Montreal, Canada, kindly provided the K27M-mutant-deleted BT245 cells, which were cultured as described previously (25). The deletion of the mutant in these cells was periodically confirmed by western blotting using K27M and K27Me3 antibodies by us.

### RNA analysis

Total RNA from tumor and normal cells with the indicated treatments was extracted using the miRNAeasy Plus kit (Qiagen) and used for the analysis of gene expression using RT-qPCR (Four replicates) and/or barcoding, followed by RNA sequencing using the Illumina Hiseq 4000 instrument.

1. *<u>Gene expression analysis:</u>* Total RNA isolated from cells with high purity was used to analyze gene expression by RT-qPCR performed on a Step One Plus Real-Time PCR System using TaqMan gene assay reagents (Applied Biosystems), according to the manufacturer’s protocols, as described by us previously (61) and using gene primers (See list in **Supplementary Table S3**). cDNA was generated from total RNA using a high-capacity cDNA kit (Applied Biosystems). The resulting cDNA was used as an input to each RT-qPCR, along with the respective TaqMan gene-specific probe for each gene set and PCR reagents. All RT-qPCR assays were performed in quadruplicate. Relative quantity was calculated using the ΔΔCT method with GAPDH as the endogenous control. Gene expressions in a larger cohort of patient samples were derived from a public dataset using the R2 Genomics Analysis and Visualization Platform (https://hgserver1.amc.nl/cgi-bin/r2/main.cgi).
2. *<u>RNA sequencing data analysis:</u>* RNA sequencing data were analyzed for differential gene expression using R. Differentially expressed genes were visualized using a volcano plot. Metascape (https://metascape.org/gp/index.html) was used to identify signaling pathways using the top 25% differentially expressed genes and are visualized using Cytoscape (https://cytoscape.org).

### Single-cell RNA sequencing (scRNAseq) and analysis

scRNAseq was performed as previously described (63). After demultiplexing, the raw sequencing reads were mapped to the human reference genome (build GRCh38), and gene-expression matrices were generated using Cell Ranger (version 3.0.1). The resulting count matrices were further filtered in Seurat 3.1.0 to remove cell barcodes with less than 200 genes (mitochondrial genes or putative doublets), yielding 18500 single cells across all samples. After normalization, based on dimensionality reduction by PCA using the 1,602 most variable genes, these cells were clustered using the Seurat workflow. Inter-sample variation due to experimental or sequencing batch effects was corrected using harmony alignment (theta = 2). After the assessment of clustering using a variety of dimensions, we used the default setting of the first 20 harmony dimensions to cluster the data and performed dimensionality reduction using Uniform Manifold Approximation and Projection (UMAP). Differential expression and marker gene identification were performed using MAST (64). Non-neoplastic cell populations were excluded, and the cells were re-clustered and re-projected to refine the neoplastic subpopulation-specific differential expression gene lists, which were further filtered to remove ribosomal protein genes. Chromosomal CNVs of single cells from the scRNaseq were inferred based on the average relative expression in variable genomic windows using InferCNV. Cells classified as non-neoplastic were used to define a baseline of normal karyotype such that their average copy number value was subtracted from all cells. Neoplastic subpopulations were characterized by direct examination of neoplastic-subpopulation-specific gene lists, enrichment of comparator gene sets, and inference of regulatory gene networks. DAVID (Database for Annotation, Visualization, and Integrated Discovery, version 6.8) was used to measure the enrichment of GO term Biological Process Direct gene sets in subpopulation signatures, providing gene ontology mappings directly annotated by the source database. Subpopulation signature gene sets included the top 250 differentially expressed subpopulation markers ranked by adjusted p-values and were filtered to remove ribosomal genes. Enrichment of selected comparator gene sets was performed using hypergeometric testing.

### Protein analysis

#### Western blot protein analysis

Protein analysis was performed with whole cell lysates isolated from DMG patient tumors and cell lines with or without the indicated treatments/modifications. Protein lysates were collected in RIPA buffer supplemented with protease inhibitor cocktail tablets, sodium vanadate, and sodium molybdate, as previously described (61). To perform histone protein analysis, histone proteins from the cells were acid-extracted using 0.2N HCL as described previously by us (61). Western blotting was performed after the protein concentrations were determined using the BCA Assay. The antibodies used for western blotting are listed in **Supplementary Table S2**.

#### Immunohistochemistry (IHC)

IHC was performed using standard protocols. Formalin-fixed, paraffin-embedded slices of primary patient tumor tissues, tissue arrays from non-cancerous specimens, and tissues isolated from mice were used for IHC. In experiments to evaluate the ability of the antibody to cross the blood-brain barrier and to reach the pons tumor site, antibody-treated mice were subjected to transcardiac profusion, and the mice brains were collected for IHC following the protocol described previously (65). The list of antibodies used to detect different proteins using IHC is provided in **Supplementary Table S2**.

#### Immunofluorescence (IF)

shRNA-mediated knockdown of CD99 was also analyzed by IF. DMG cells (∼30,000 cells/well) were cultured on poly-d-lysine/laminin 8-well culture slides (n = 2 per cell line). The next day, the treatment medium was aspirated, and the cells were fixed with 4% paraformaldehyde for 15 minutes at room temperature, followed by washing twice with 0.2% Triton-X-PBS. Cells were incubated in the blocking solution of 5% milk in 0.05% Triton-X-PBS on a shaker for 30 minutes at room temperature, then incubated with the primary antibody overnight at 4°C. The following day, the cells were washed twice with 0.05% Triton-X-PBS and then incubated with the secondary antibody to CD99 (Alexa Fluor 488, Green) and GFAP (Alexa Fluor 568, Red) in 5% milk in 0.05% Triton-X-PBS for 1 hour at room temperature in the dark. Complete details of the primary and secondary antibodies are provided in **Supplementary Table S2**. The slides were washed with 0.05% Triton-X-PBS, washed three times with 1X PBS, allowed to air dry for 3 min, and the coverslip was mounted with ProLong™ Gold Antifade Mounting reagent containing DAPI (Invitrogen) to label the nuclei. The slides were then imaged for each protein using a Keyence microscope. The expression of CD99 and GFAP was analyzed using the Hybrid Cell Count plugin in the BZ-X Analyzer Software.

### Flow Cytometry

#### Cell Surface Expression of Proteins

The cell surface proteins were detected, and their expression was quantitated using a CytoFlex LX flow cytometer. Briefly, single-cell suspensions of unmodified or modified DMG and normal cells were spun at 300g for 5 min and resuspended in cell staining buffer containing conjugated (given in **Supplementary Table S2**) or unconjugated (10D1) primary antibodies. The cells were then incubated in the dark at 4°C for 30 minutes. After the incubation, the cells were washed twice with cell staining buffer and resuspended in the corresponding anti-secondary antibodies. 30 minutes after this incubation in the dark at 4°C, the cells were washed twice and resuspended in cell staining buffer containing 0.1μg/ml DAPI. The cells were then transferred to a Cytoflex LX flow cytometer to record the mean fluorescence intensity and the percentage of DAPI-negative live cells. All samples were run on a flow cytometer and the data obtained were analyzed using FlowJo® software. The results are represented as a histogram of the cell count of mean fluorescence intensity, representing the size of the DAPI-negative live cell population emitting fluorescence, and a bar graph representing the mean fluorescence intensity of specific proteins across different cell lines.

#### Antibody blocking experiments

DMG cells were treated with either IgG4 or with varying concentrations between 10-50μg of 10D1 antibody first with the antibody suspended in 1X PBS at 4°C for 30 minutes. After aspirating the PBS, the cells were incubated again with the antibody in serum-free media at 37°C for 4 hours. The extent of CD99 blocking by the 10D1 antibody was measured using flow cytometry with CD99-0662 as the detection antibody. Stained cells were washed and resuspended in cell staining buffer containing 0.1μg/ml DAPI, and the mean fluorescence intensity was recorded in the percentage of CD99-0662^+^ cells amongst the DAPI-negative live cells on a Cytoflex LX flow cytometer and analyzed as described above.

### ChIP and ChIP-Sequencing (ChIP-seq)

Chip-seq was performed as previously described (61). Briefly, cells with or without the indicated gene modifications were fixed with 1% formaldehyde in 1X DPBS for 10 minutes at RT. The cells were washed twice in 1X PBS and resuspended in RIPA buffer with a protease inhibitor cocktail. The cell lysates were then sonicated in a Diagenode sonicator to yield chromatin fragments with DNA of size 250-500bp, verified using an agarose gel electrophoresis assay. BCA assay was performed on the chromatin fragment after the removal of debris. 1mg of chromatin fragments were then immunoprecipitated at 4°C for 1 hour with 40μg of protein A/G agarose beads to pre-clear with RIPA buffer. The beads were centrifuged and pelleted, and the supernatant was collected in a fresh tube. Pre-cleared chromatin extracts were incubated with 2μg of primary antibody and fresh protein A/G agarose beads at 4°C overnight with slow rotation. The immune precipitates were washed successively with 1 mL of low salt buffer (20mM Tris-HCl [pH 8.0], 150mM NaCl, 0.1% SDS, 1% Triton X-100, 2mM EDTA), high salt buffer (20mM Tris-HCl [pH 8.0], 500mM NaCl, 0.1% SDS, 1% Triton X-100, 2mM EDTA), LiCl washing buffer (10mM Tris-HCl [pH 8.0], 250mM LiCl, 1.0% NP40, 1.0% deoxycholate, 1 mM EDTA) and twice with TE buffer. The DNA-protein conjugated with the antibody complexes was eluted with 200μl of an immunoprecipitation elution buffer containing 1% SDS and 0.1 M NaHCO_3_. After pooling the eluents together and the crosslinking was reversed with 0.2M NaCl, they were incubated for 5 hours at 65°C. DNA recovery was performed by digestion with proteinase K and RNAse A, followed by extraction with phenol/chloroform and precipitation in 200-proof ethanol. The pellets were resuspended in 100μl of 10 mM Tris-HCl (pH 8.0), and the resulting DNAs were sequenced at the Genomics Core at the University of Colorado Anschutz Medical Campus.

### Neurosphere assay

DMG cells (SU-DIPG04, BT245, and SU-DIPGXIII) with CD99 shRNA-mediated knockdown or CRISPR knockout and non-target control were seeded (∼1000 cells/well) in a 96-well low-attachment round bottom plate on day 0 (Four replicates). In the second experiment, DMG cells (BT245 and MAF-002) were seeded (∼1000 cells/well) in a 96-well round-bottom plate on day 0. On day 1, the cells were treated with either IgG4, 10μg or 25μg 10D1 antibody. They were then imaged on an Incucyte S3 Live Cell Analysis System at regular time intervals, and the change in the size of the neurosphere (i.e., the size of the largest bright field object) was analyzed and plotted against time.

### Cell proliferation assay

#### XCELLigence cell growth assay

SU-DIPG04 cells were seeded (∼10,000 cells/well) in a gold-plated 96-well E-plate, and the cell growth index was measured in real-time using xCELLigence on day 0 (Four replicates). On day 1, the cells were treated with either IgG4 or 10D1 antibodies, and the changes in the cell growth index were measured over time.

#### Incucyte cell proliferation assay

DMG cells (BT245 and MAF-001) were seeded (∼2000 cells/well) in a 96-well flat bottom plate on day 0 (Four replicates). On day 1, the adherent cells were treated with either IgG4 or 10D1 antibody. They were then imaged on an Incucyte S3 Live Cell Analysis System at regular time intervals, and the percentage of phase confluence as a measure of the number of viable cells was analyzed and plotted against time.

### ALDEFLUOR^TM^ Assay

ALDEFLUOR^TM^ Assay was performed based on the manufacturer’s protocol (Stemcell Technologies #01700). Briefly, BODIPY-aminoacetaldehyde-diethyl acetate or simply ALDEFLUOR™ reagent powder was dissolved in DMSO and treated with 2N HCl to convert it into fluorescent-activated ALDEFLUOR™ reagent, diluted with ALDEFLUOR™ assay buffer, and maintained at 4°C. CD99 knockdown SU-DIPG04 cells and non-target controls were suspended in ALDEFLUOR™ assay buffer at a concentration of 1 × 10^6^ cells/mL in two 1.5mL Eppendorf tubes for each cell line (Three replicates). One of the sample tubes was treated with 5μL of ALDEFLUOR™ Diethylaminobenzaldehyde reagent (control), and the other tube was treated with 5μL of ALDEFLUOR™ reagent. After mixing the reagents in each tube, 500μL of the control tube suspension was added to the test tube and mixed thoroughly again. Control and substrate buffers were then added to the respective control and test tubes, and the cells were incubated at 37°C for 30 min. After incubation, the cells were pelleted (300g for 5 minutes) and resuspended in the ALDEFLUOR™ assay buffer. The fluorescence activity in the stained control and test samples was evaluated on a Guava Flow Cytometer, and the percentage of ALDH^+^ cells, defined as those emitting fluorescence higher than the control, was quantified using FlowJo ® software.

### Co-immunoprecipitation

To determine whether the two proteins, CD99 and ILK1 interacted directly or indirectly, Co-Immunoprecipitation (Co-IP) was performed using the Co-IP kit (Active Motif) according to the manufacturer’s instructions and as described previously (33). Briefly, DMG cells (SU-DIPG04, HSJD-DIPG007, SU-DIPGXIII) were collected in 1X PBS supplemented with phosphatase and deacetylase inhibitors, and the whole-cell lysates were isolated using the Complete Whole-Cell Lysis buffer. 500 μg of whole cell lysate was incubated with 2.5μg of ILK1 or CD99 antibody for 4 hours at 4°C on a shaker. The whole cell lysate-antibody conjugates were incubated with 25μL of Protein G Magnetic beads for 1hr at 4°C on a shaker. After incubation, conjugates were collected using a magnetic stand and washed. The interaction of CD99 with ILK1 was evaluated by western blotting of proteins immunoprecipitated with either CD99 or ILK1 antibodies (See **Supplementary Table S2**).

### Drug dosing and IC_50_ determination

DMG Cells (∼2,000 cells/well) were seeded on a flat-bottom 96-well plate (SU-DIPG04, HSJD-DIPG007) or round-bottom low attachment 96-well plate (BT245, SU-DIPGXIII) with and without gene-modifications on day 0 (Four replicates). On day 1, the DMG cells were treated with a range of concentrations of cpd22 (0-2μM). Concentration-matched DMSO-treated cells were used as controls. On day 4, an MTS assay was performed to determine changes in the cell viability. The non-linear fit analysis of the drug dosing data (absorbance) was performed on the GraphPad Prism software ® to determine the amount of cpd22 needed for the death of 50% of the cells (IC_50_) in DMG cells.

### *In vitro* irradiation

DMG cells were grown to 70% confluence and were exposed to either a single irradiation dose (2, 4, or 6 Gy) or fractionated doses of radiation (2 Gy/day for three days or 4 Gy/day for three days) using an X-ray irradiator. Non-radiated cells were used as controls. All irradiation was performed using a perpendicular 662 keV g-photon beam with dose coverage to uniformly irradiate all samples at a dose rate of 1.09 Gy/min and a source-to-surface distance of 30 cm. Irradiated and corresponding non-irradiated cells were used for the following experiments.

1. To determine the effect of the combination of radiation and antibody treatment, 24 hrs after the final dose of radiation, cells were treated with varying concentrations of either control or anti-CD99 (10D1) antibody (Three replicates). At 24, 48, and 72 hrs after incubation with the antibody cell apoptosis was measured using Incucyte S3 Live Cell Imaging as described in the Cell apoptosis using Incucyte section.
2. To determine the changes in the expression of CD99 after radiation treatment, 24 hrs after the final dose of radiation, cells were washed, collected by scraping, and examined by flow cytometric analysis for changes in the expression of CD99, as described in the flow cytometry section.

### Cell apoptosis measurement using Incucyte

To evaluate the induction of apoptosis by the 10D1 antibody on DMG cells, approximately 2000 cells/well were cultured in a flat-bottom 96-well plate coated with GelTrex ® (SU-DIPG04 and BT245) or a low-attachment round-bottom plate (SU-DIPGXIII) on day 0 (three replicates). On day 1, the cells were treated with 10μg of 10D1 or IgG4 antibody. Two hours after this treatment, the CellEvent Caspase3/7 Green Detection Reagent (Invitrogen) was added to each well at a final concentration of 2μM. Apoptosis induced by the indicated treatments was measured using an Incucyte S3 Live Cell Analysis System. An increase in green fluorescence indicated an increase in apoptosis over time, as measured by the amount of dye cleaved by the activation of caspase 3/7. The number of green objects was counted and expressed as Total Green Object Integrated Intensity (GCUxμm²/well), which was normalized to the percentage cell confluence, i.e., phase confluence, calculated from the phase contrast image of that well at the corresponding time point. The change in normalized intensity was plotted against time. The resultant changes in apoptosis at 24 and 48hrs of treatment with the 10D1 antibody are represented as histograms. For studies involving the 10D1 antibody without radiation, the increase in apoptosis in cells treated with the 10D1 antibody was compared with that in IgG4-treated control cells.

### *In vivo* xenograft models

Male and Female NSG mice (#005557) and female nude mice (#007850 JNU) 6-8 weeks old, were purchased from Jackson Laboratory. Mice were housed in OLAR-accredited animal facilities at the University of Colorado AMC under the University of Colorado Institutional Animal Care and Use Committee (IACUC) by the Office of Laboratory Animal Resources (OLAR) at UC AMC protocol approved. All *in vivo* mouse DMG xenograft studies, including implantation, animal care, radiation, treatments, and euthanasia, were performed under an approved animal protocol.

#### DMG tumor engraftments

Human DMG cells tagged with luciferase-GFP; BT245-Luc2-GFP, MAF-001-Luc2-GFP, and SU-DIPGXIII*-Luc2-GFP were implanted in the pons of NSG mice. BT245-shNull-Luc2-GFP and BT245-shCD99-Luc2-GFP were implanted in the pons of nude mice. The luciferase-GFP tagged 1×10^5^ cells in 2µl serum-free media were stereotactically injected at a rate of 600nL/minute into the brain at a site 0.8 mm lateral to the midline, 0.5 mm posterior to lambda, and 5.00 mm ventral to the surface of the skull (27). Tumor formation was monitored by bioluminescence imaging (BLI) once per week using a PerkinElmer IVIS Spectrum *in vivo* imaging machine.

In shRNA-mediated-CD99 knockdown xenografts, tumor growth and body weight were monitored during the study. Mice were monitored daily for any signs of lethargy, such as weight loss, hunched position, and spin. Tumor growth was monitored weekly using IVIS. At the end of this study, the mice were euthanized, and the tumors and brains were collected for total RNA extraction and IHC.

#### Antibody treatment

The tumor take rate was 100%. After confirmation of tumor engraftment in pons with BLI corresponding to approximately 10^4^ to 10^6^ radiance (p/sec/cm^2^/sr) using IVIS, the mice were randomized for different treatments as follows.

Two groups: group 1, cohort of mice that received IgG4 control, and group 2, cohort of mice that received 10D1 (anti-CD99) antibody.

#### Intravenous delivery of antibodies

The control or anti-CD99 (10D1) recombinant antibodies were infused *via* tail vein injection (IV) in the pons tumor-bearing mice, at a dose of 8 mg/kg of body weight resuspended in 200 µL volume of sterile ice-cold 1X PBS for consecutive days after tumor inoculation, and changes in tumor growth were monitored.

#### Loco-regional delivery of antibodies to CNS

Antibodies at a single dose of 1 mg/kg of body weight in 5μl total volume were stereotactically injected at a rate of 600nL/minute into the lateral ventricular area of mice intrathecally (IT) using the following co-ordinates with respect to Bregma 1.0 mm lateral to midline, -0.5 mm anterior to lambda, and -3.00 mm ventral to the surface of the skull and changes in tumor burden was monitored.

#### Antibody in combination with radiation treatment (RT)

Two sets of experiments were performed to understand the effect of blocking CD99 in combination with RT on tumor growth. In the first set of experiments, antibodies were delivered by IV; in the second set of experiments, antibodies were delivered by IT injection. In both sets of experiments, antibodies were delivered using the exact coordinates described above. After the orthotopic DMG model was established, for each of the two sets of experiments, mice were randomized into the following treatment groups: (i) control, (ii) RT only, and (iii) antibody followed by RT (2Gy/day × 3). Image-guided irradiation was delivered using an X-RAD SmART small-animal irradiator (Precision X-Ray, Madison CT), using a beam of 225 kVp and 20mA, with a 6.01 Gy/min dose rate. Mice were anesthetized and placed in the prone position on the irradiator bed, with their nose in a nosecone delivering in-machine anesthesia. Fluoroscopy was used to center the pons at the isocenter in two orthogonal planes. Half of the prescribed dose per fraction (2Gy) was delivered to either side of the head to create a more uniform dose distribution, which was confirmed using a Monte Carlo simulation using SmART-ATP treatment planning software (SmART Scientific Solutions, Maastricht, Netherlands). Irradiation using the above procedure was repeated for three days, delivering three fractions of 2Gy to the pons.

Mice were monitored daily for signs of lethargy, such as weight loss, hunched position, head tilt, and spin. Tumor growth and antibody therapy responses were determined weekly using BLI. Tumor volume was calculated from the total flux intensities (66). The total photon flux in each mouse was determined using a specific region of interest in animals and integrating photon flux over the entire imaging period. Mice body weight was measured once a week, and those reaching the endpoint were euthanized according to IACUC protocols by CO_2_ asphyxiation when they showed signs of either neurologic deficit, head tilt, failure to ambulate, body score less than 2, or weight loss greater than 20%.

At the endpoint, mice were anesthetized, and blood samples were withdrawn by heart puncture and collected in EDTA-coated tubes to check if any toxicity occurred after antibody treatment, either alone or in combination with radiation treatment. Animals were anesthetized for exsanguination. Hematological parameters were measured in the collected blood using a complete blood count (CBC) instrument, Element HT5 Veterinary Hematology Analyzer (Heska; n=5, biological replicates per treatment group). After collecting blood samples, the mice were euthanized, and residual tumor tissues and brains were collected for IHC analysis.

#### Magnetic Resonance Imaging (MRI) of the mouse brain

High-resolution T2-turboRARE MRI of both sagittal and axial planes was acquired on a Bruker 9.4 Tesla BioSpec MRI scanner using a mouse head array coil to visualize changes in tumor volume. Tumor volumes (mm^3^) were calculated by placing a region of interest (ROI) on each anatomical slice and multiplying the sum by the slice thickness (0.7 mm). Tumor volume analysis was performed by radiologists blinded to the treatment groups.

#### Chimeric anti-CD99 (10D1) antibody development

The novel chimeric anti-CD99 antibody was generated as human IgG4 molecules under contract with GenScript (Piscataway, MJ). Genscript synthesized linear peptides with sequences corresponding to amino acids 75-85 for the CD99 antigen. Antisera from mice inoculated with this peptide were used for screening using an immunoprecipitation assay. We performed an immunoprecipitation assay with antisera to determine the specific binding to CD99 and identified the mice with the most efficient IP in DMG. GenScript then used these selected mice for thymus cell fusion to produce hybridoma clones. We tested the supernatants from the different hybridoma clones for binding affinity, blocking ability, and tumor cell lysis function and selected the most potent clone, 10D1. To measure the kinetics of the 10D1 antibody association and dissociation, surface interferometry (Octet Red 384) was used. Octet system Data Analysis software was used to create kinetic curves to equations based on 1:1 kinetics and calculate KD, Ka, and Kd(off).

#### Experimental design and statistical rationale

##### *In vitro* assays

Most *in vitro* assays were repeated at least twice. RNA sequencing was performed. Student t-tests and two-way ANOVA were used for comparison.

##### *In vivo* assays

Animal studies were performed in cell lines and PDX models to strengthen experimental rigor. The mouse experiments had equal numbers of male and female mice evenly divided in each treatment group. ANOVA statistical analysis with appropriate post hoc tests was used for multiple comparisons. P-values were adjusted for multiple testing using Bonferroni correction. Using the Kaplan-Meier method, a survival curve was created based on the time to the endpoint value, and groups were compared using the log-rank Mantel-Cox test.

Each figure or legend indicates the statistical p-value used. For all figures, * indicates p<0.05, **p<0.01, ***p<0.001, ****p<0.0001, and ns indicates non-significant, p>0.05. Graphs were created using GraphPad Prism software. All error bars indicate standard errors of the mean (SEM) except where specified.

## Acknowledgments

The authors appreciate the contribution made by the University of Colorado Denver Tissue Histology Shared Resource, the Genomics Core, and the Small Animal Irradiator Shared Resource, all supported in part by the Cancer Center Support Grant (P30CA046934). The authors thank Jenna Steiner and Dr. Natalie Serkova for assistance with the MRI imaging (CU Cancer Center Animal Imaging Shared Resource). We acknowledge the services provided by the Pathology Shared Resource and the Molecular Biology Core Facility at the UCD Barbara Davis Center for performing mycoplasma testing and fingerprinting of the various cell lines. The authors thank Etienne Dannis for consultation on informatics. The study was supported by the Luke’s Posse Fund (SV, RV), Morgan Adams Foundation (SV, RV), The Cancer League of Colorado (SV, RV), Chad Tough Foundation (SV), Olivia Caldwell Foundation (SV), Marc Jr. Foundation (SV), Laya Dance Arts (SV), the Department of Defense (CA190645, SV), and the CU Innovations/NIH REACH-SPARK Award (RV, SV).

## Author Contributions

IB, KM, and SV contributed to the design and implementation of this study. IB, KM, DW, JM, ND, SLC, JD, BB, ZN, KJ, ND, AG, AP, and AD contributed to the development and implementation of the methodology. IB, KM, SV, and RV analyzed the results. IB, KM, SV, and RV wrote and edited the manuscript. SV and RV conceived and supervised the study.

Conflict of Interest declaration: IB, KM, DW, JM, ND, SLC, JD, BB, ZN, KJ, ND, AG, AP, and AD report no affiliations with or involvement in any organization or entity with any financial interest in the subject matter or materials discussed in this manuscript. SV and RV are patent holders for 10D1 and co-founders of Vinasa Oncology.

## Supplementary Figures

**Figure S1:**
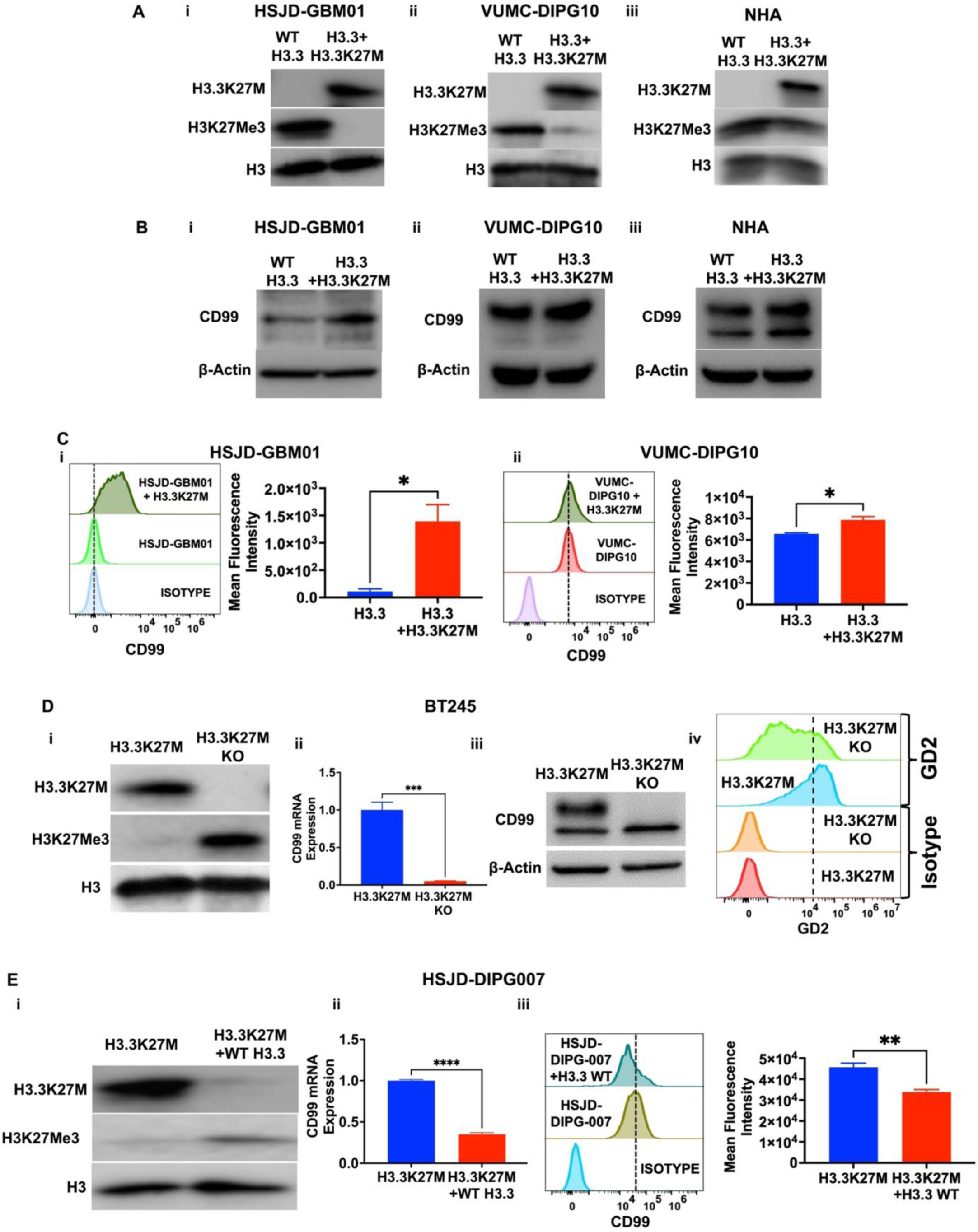
Altering somatic (K27M) mutations of the genes encoding the histone H3 variants in neoplastic cells alter the expression of CD99. A. DMG and normal cells were transduced with K27M mutant at the H3.3 locus. Western blot showing the changes in histone H3.3K27M and histone H3K27Me3 proteins in the transduced cells of (i) HSJD-GBM01, (ii) VUMC-DIPG10, and (iii) NHA, respectively. B. In the above-transduced cells, the changes in the expression of CD99 in (i) HSJD-GBM01, (ii) VUMC-DIPG10, and (iii) NHA, respectively are shown by western blotting. C. Flow cytometry plots for CD99 (i) HSJD-GMB01 transduced with H3.3K27M, and (ii) VUMC-DIPG10 transduced with H3.3K27M. The quantification of the mean fluorescence intensity is shown adjacent to the flow plots. (*p<0.05) D. H3.3K27M mutant in BT245 cells was deleted by Crispr-CAS9 gene editing. (i) Expression of histone H3.3K27M and histone H3K27Me3 in the paired samples by western blotting. (ii) H3.3K27M mRNA expression by RT-qPCR. (iii) Expression of CD99 in the paired cell lines by western blotting. (iv) Expression of the cell surface marker, GD2, by flow cytometry. (***p<0.001) E. HSJD-DIPG007 cells were transduced with H3.3 Wildtype using shRNA. (i) Expression of histone H3.3K27M and histone H3K27Me3 in the paired samples by western blotting. (ii) H3.3K27M mRNA expression by RT-qPCR. (iii) Expression of CD99 in the paired cell lines by western blotting. (**p<0.01, ****p<0.0001) The expression of total histone H3 or β-Actin was used as protein loading control in all western blots. IgG, respective to each antibody, was used for gating cells in flow cytometry. DAPI was used to gate for live cells. GAPDH was used as a control for qRT-PCR. Data represented as Mean ± SEM.

**Figure S2:**
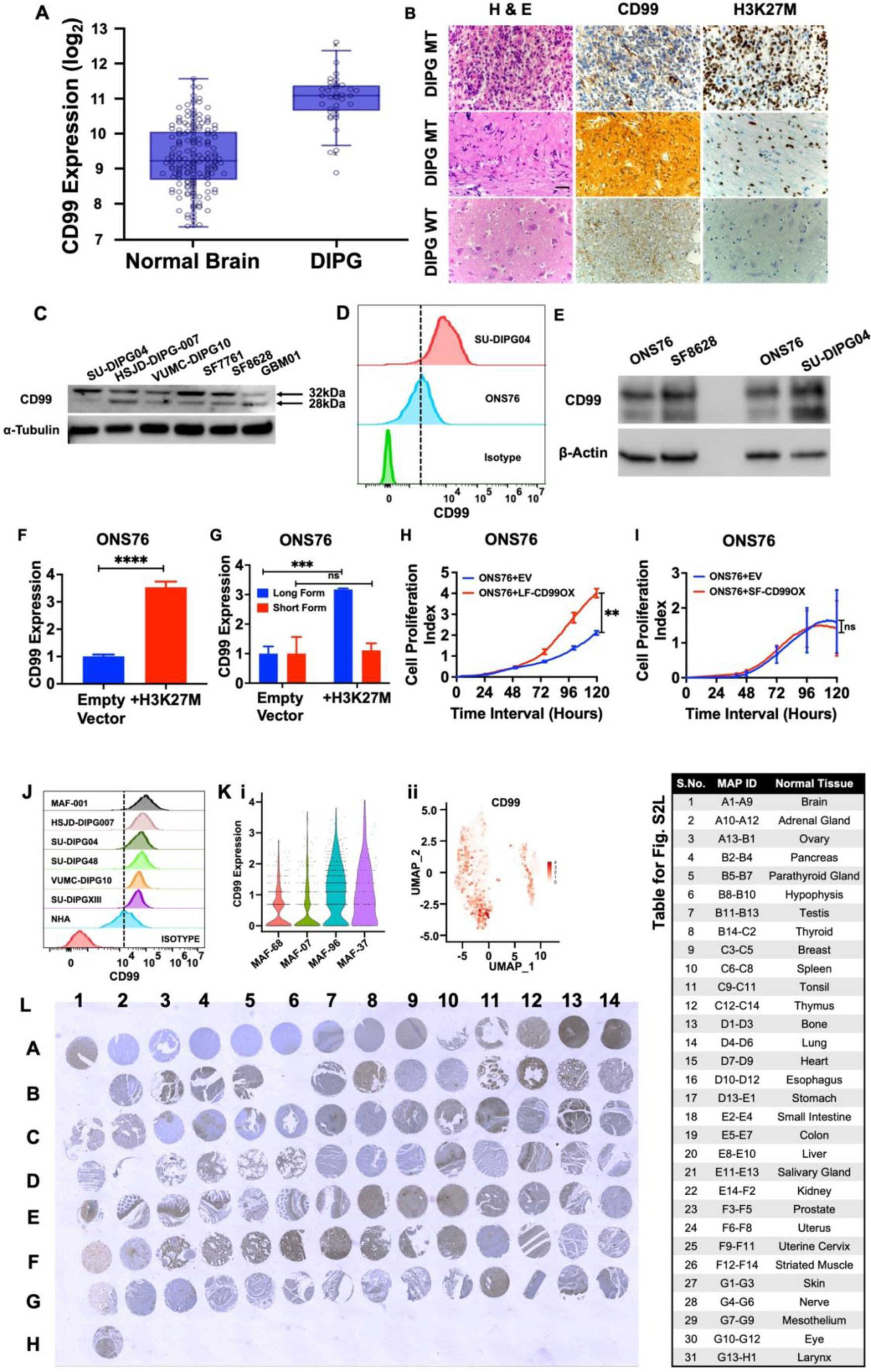
CD99 and the role of its isoforms in DMG proliferation. A. CD99 expression in a large cohort of DIPG patient samples (n=37) (67) compared to its expression in normal brain regions (n=172) (68) from the dataset using the R2 genomics. (Data represented as mean ± SD.*p<0.0001) B. Representative IHC staining showing expression of CD99 in H3.3 wildtype and H3.3K27M mutant DMG tumors. The mutation status was verified using H3K27M staining. C. CD99 protein expression in cultured DMG cell lines is shown by western blotting. D. The sonic hedgehog-activated medulloblastoma line, ONS76, has a relatively lower expression of CD99 compared to the K27M mutant DMG cell line, SU-DIPG04 as seen by flow cytometry. E. ONS76 demonstrates low expression of CD99 compared to the K27M mutant DMG cell lines, SF8628 and SU-DIPG04 as seen by western blotting. F. The overexpression of the K27M transgene in ONS76 was verified using RT-qPCR. (****p<0.0001) G. The overexpression of the K27M transgene in ONS76 significantly upregulated the long isoform of CD99 but not the short isoform by using the respective primers. (ns = not significant, ***p<0.001) H. Overexpression of the long isoform of CD99 transgene significantly enhanced the proliferation of ONS76 cells. (**p<0.01) I. Overexpression of the short isoform of CD99 transgene did not alter the proliferation rate of ONS76 cells. (ns = not significant) J. Expression of CD99 by flow cytometry in cultured DMG cell lines including (i) DMG H3 wild type lines, VUMC-DIPG10 and SU-DIPG48, (ii) H3K27M mutant lines, SU-DIPGXIII, SU-DIPG04, HSJD-DIPG007, (iii) a primary cell line, MAF-001, and (iv) NHA. K. (i) Violin-plot showing the expression of CD99 transcripts at single cell level in each of the four DMG tumor cohorts collected at the Children’s Hospital, Colorado. (ii) Corresponding Uniform Manifold Approximation and Projection (UMAP) plot showing the expression of CD99 transcript at single cell level. L. IHC for Normal human Tissue Array (FDA999-1) stained for the expression of CD99 and the adjacent table gives a brief description of the tissues in the array collected from respective organs (Refer to https://usbiolab.com/tissue-array/product/multiple-organ/FDA999-1 for the list of tissues on this array and absence of the H&E staining) Data represented as Mean ± SEM.

**Figure S3:**
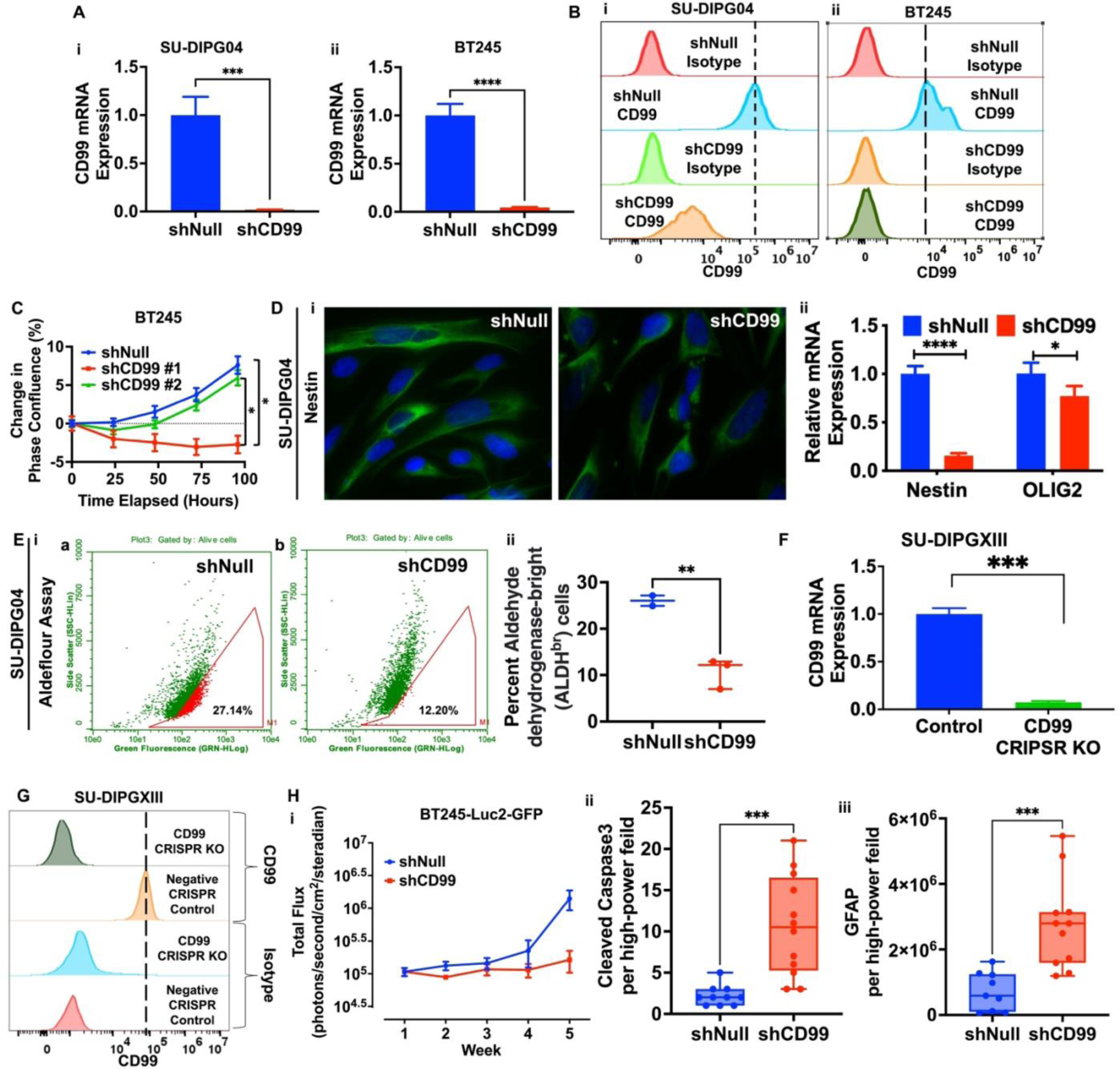
Knockdown or deletion of CD99 impairs DMG growth and enhances DMG cell differentiation. A. CD99 mRNA expression in shRNA-mediated knockdown of CD99 in (i) SU-DIPG04 and (ii) BT245 cells. B. CD99 protein expression in the above cells by flow cytometry. C. shRNA-mediated knockdown of CD99 in BT245 cells, with different constructs, slows cell proliferation as measured as phase confluency using incucyte. (*p<0.05) D. shRNA-mediated knockdown of CD99 enhances the differentiation in SU-DIPG04 cells. (i) IF images of (a) control shNull and (b) shCD99 cells. The nuclei are stained with DAPI (blue) and Nestin (green). Scale bar = 50μm. (ii) mRNA expression of Nestin and Olig2 in SU-DIPG04 after shRNA-mediated knockdown of CD99 by RT-qPCR. (*p<0.05, ****p<0.0001) E. Self-renewal and differentiation of SU-DIPG04 increased after the shRNA-mediated knockdown of CD99 as measured by the Aldefluor assay. (i) Representative flow plots. (ii) Quantification of the mean fluorescence intensity of the percent aldehyde dehydrogenase-bright cells (n=3). (**p<0.01) F. CD99 mRNA expression in Crispr-CAS9 deletion of CD99 in SU-DIPGXIII cells by RT-qPCR. (***p<0.001) G. CD99 expression in Crispr-CAS9 deletion of CD99 in SU-DIPGXIII cells by flow cytometry. H. Establishment of the CD99-sufficient (shNull) and CD99-deficient (shCD99) tumors in mouse xenograft models. (i) Bioluminescence as log_10_(Total Flux) values (photons/second/cm^2^/steradian) showing changes in tumor burden in the CD99-sufficient and CD99-deficient tumor groups. (ii) Cleaved caspase3 and (iii) GFAP quantified from n>5 different high-power fields from the stained IHC regions obtained in Figure 2G(v) from n=2 mice for each group. (***p<0.001) Data represented as Mean ± SEM.

**Figure S4:**
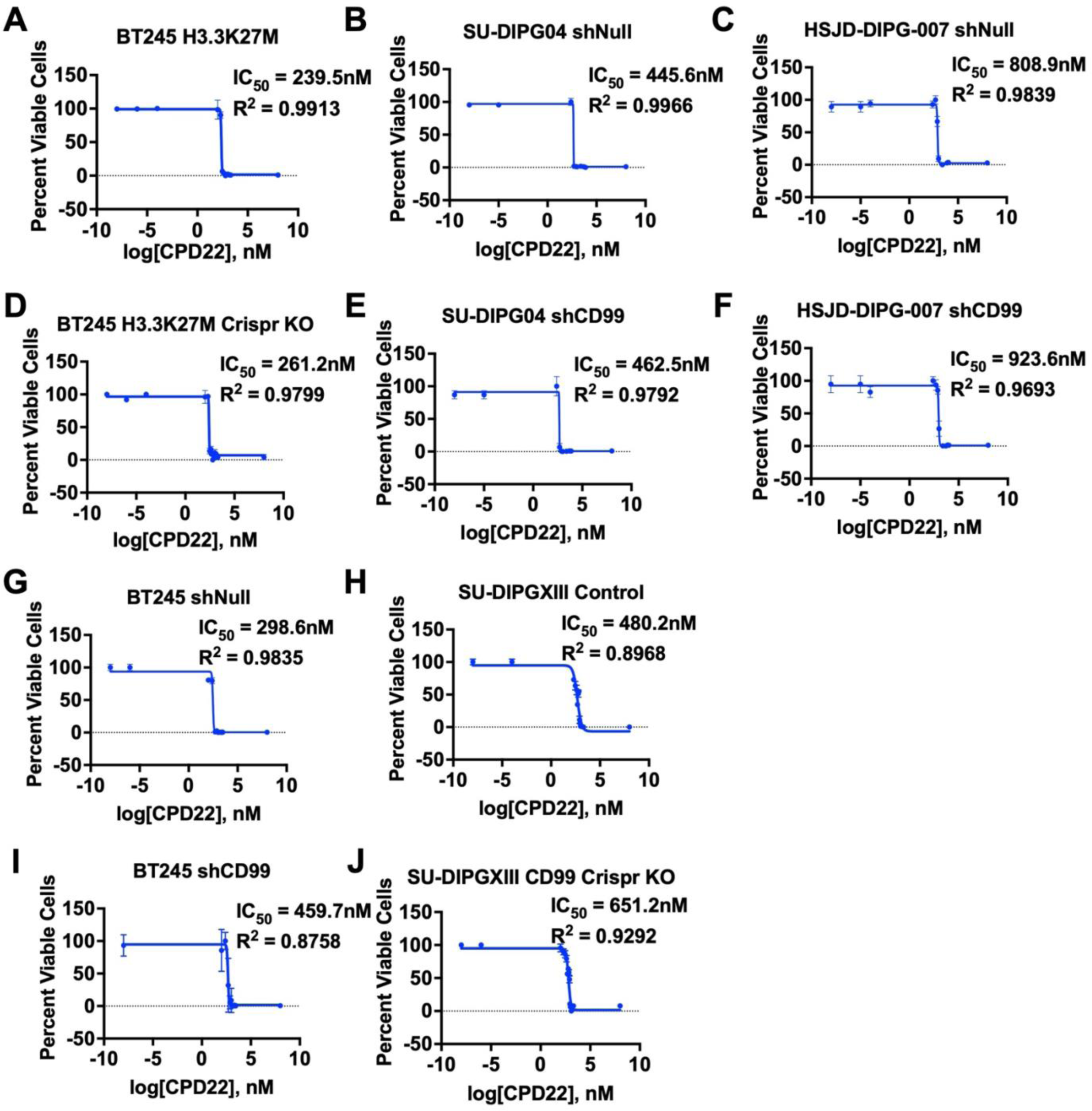
Dosing curves of cpd22 for DMG cells representing the percent viable cells against the log concentration of cpd22. A. Dosing curve for BT245 H3.3K27M parental cell line. B. Dosing curve for SU-DIPG04 shNull (non-target control) cell line. C. Dosing curve for HSJD-DIPG007 shNull (non-target control) cell line. D. Dosing curve for BT245 H3.3K27M with Crispr-CAS9 deletion of K27M mutation cell line. E. Dosing curve for SU-DIPG04 shCD99 cell line. F. Dosing curve for HSJD-DIPG007 shCD99 cell line. G. Dosing curve for BT245 shNull (non-target control) cell line. H. Dosing curve for SU-DIPGXIII parental (control) cell line. I. Dosing curve for BT245 shCD99 cell line. J. Dosing curve for SU-DIPGXIII with Crispr-CAS9 deletion of CD99 cell line.

**Figure S5:**
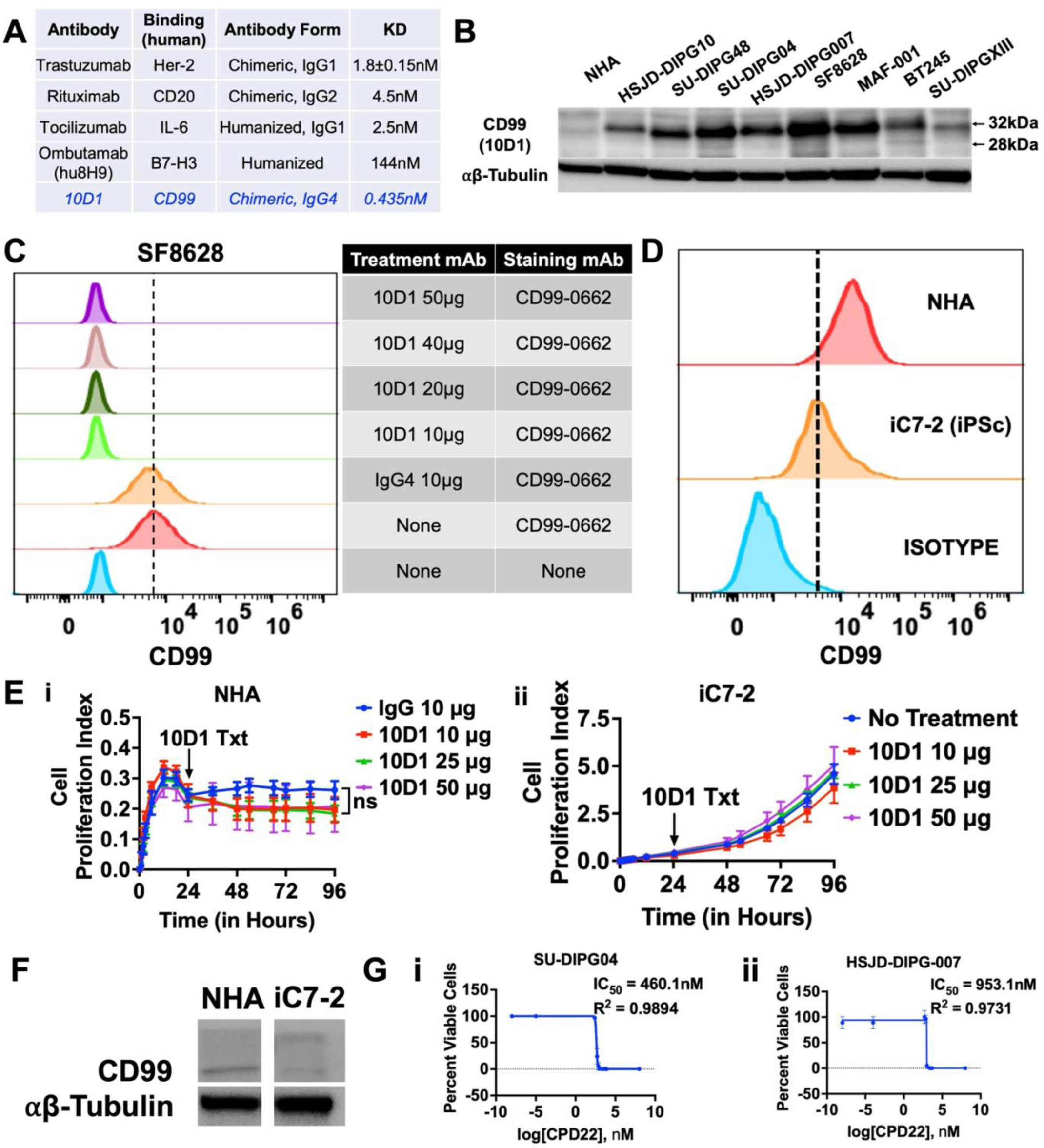
Binding affinity of 10D1 to human CD99 and efficacy of 10D1 in DMG tumor and normal cells. A. Binding affinity of clinically relevant antibodies and 10D1. B. 10D1 selectively binds to the long isoform (32kDa) of CD99 in DMG and normal cells. C. Blocking of CD99 in SF8628 using the 10D1 antibody followed by the detection with CD99-0662 antibody shows that 10D1 effectively blocks CD99 even at low concentrations. D. CD99 protein expression in NHA and induced pluripotent stem cells (iC7-2) by flow cytometry. E. 10D1, at different concentrations, did not inhibit the growth of (i) NHA and (ii) iC7-2 cells measured using xCELLigence. F. CD99 protein expression in NHA and iC7-2 by western blotting. G. Dosing curve of cpd22 representing the percent viable cells against the log concentration of cpd22 for (i) SU-DIPG04 and (ii) HSJD-DIPG007 cell lines.

**Figure S6:**
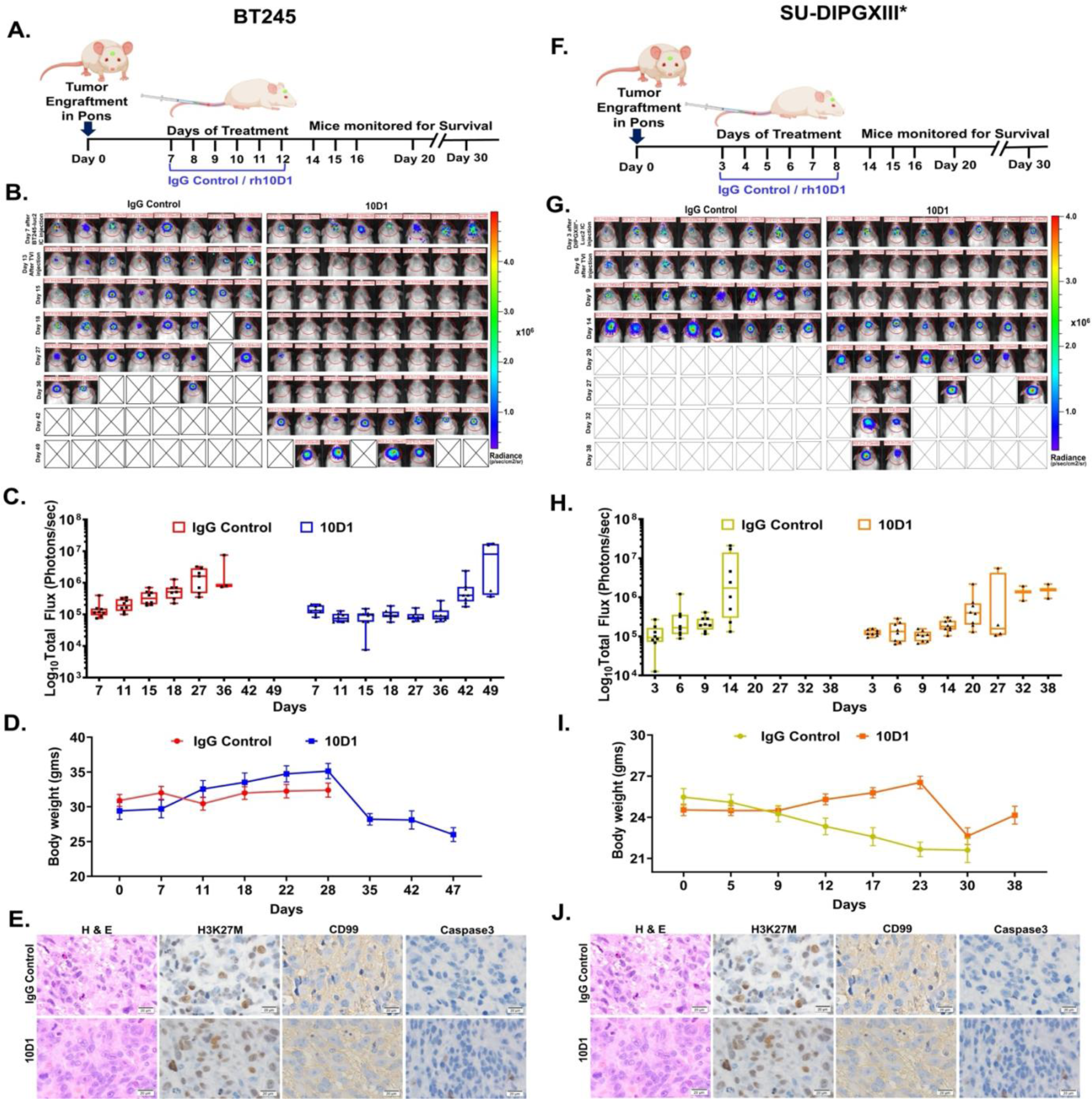
IV delivery of 10D1 restricts the DMG growth and improves survival in DMG xenograft-bearing mice. Schematic diagram of the study protocol for **A**: BT245 and **F**: SU-DIPGXIII* tumor model and treatment. **B & G**: Individual BLI of mice in each treatment cohort for different days. **C & H**: Analysis of BLI for the 10D1 and IgG treatment groups. **D & I**: Relative body weight from each treatment cohort over time. **E & J**: IHC analysis of the brain tissue collected from mice at the endpoint of the study and stained for indicated proteins. Data represented as the mean ± SEM.

**Figure S7:**
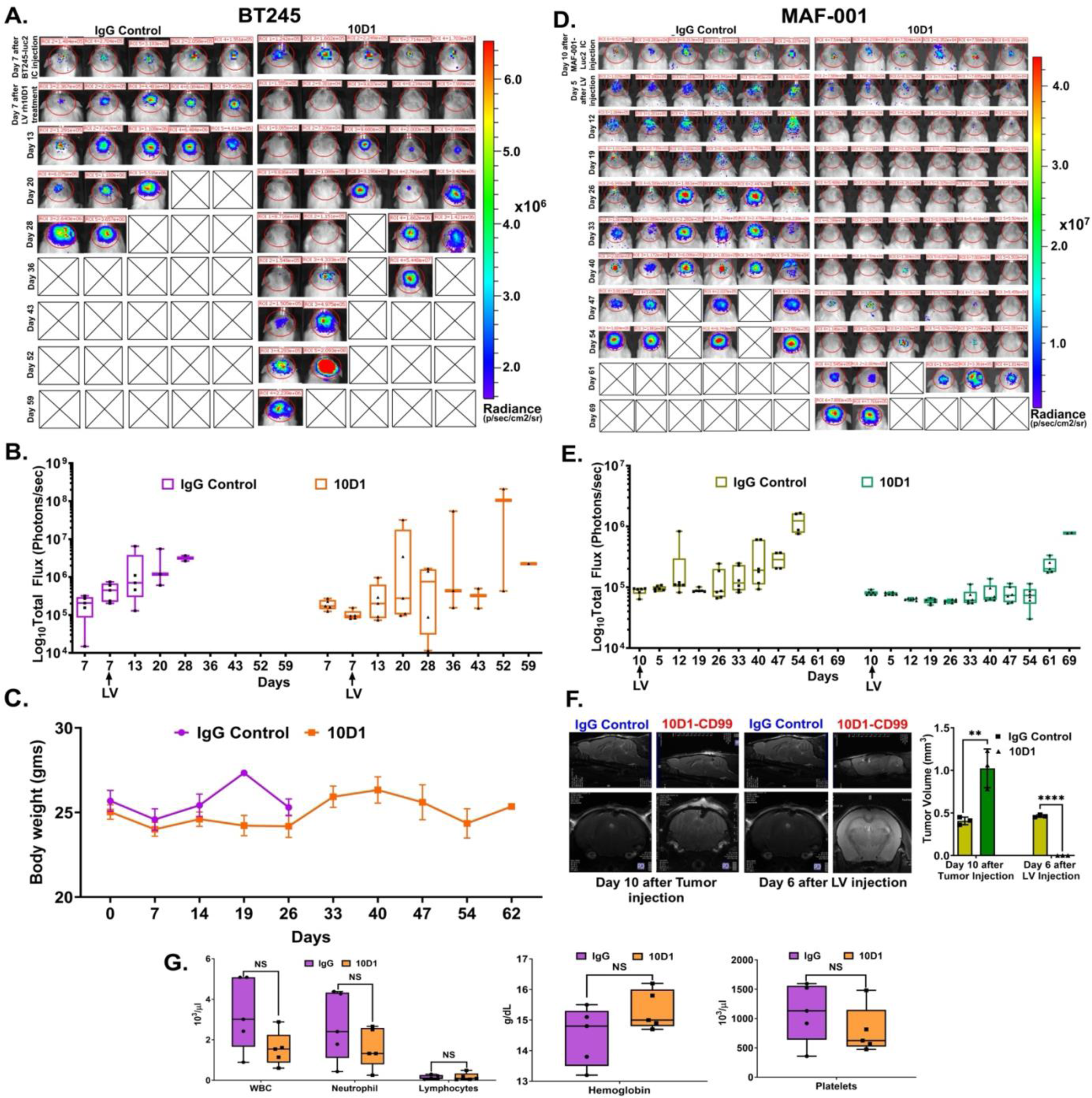
IT delivery of 10D1 inhibits DMG tumor growth in xenografts. **A**: BT245 and **D:** MAF-001 tumor models and their individual BLI of mice in each treatment cohort for different days. **B & E**: Analysis of BLI for the 10D1 and IgG treatment groups. **C**: Relative body weight of BT245 model in both groups. **F:** MRI imaging of the MAF-001 model for IgG control and 10D1 treatment showing tumor clearance and its quantitative analysis of tumor volume significantly reduced at day 6 after a single dose of IT delivery (****p<0.001). **G**: Complete blood count (CBC) analysis from the blood collected immediately after animals were euthanized upon reaching their endpoint. Samples were analyzed for WBC (leukocytes), hemoglobin, neutrophils, lymphocytes, and platelets count. Data represented as the mean ± SEM.

**Figure S8:**
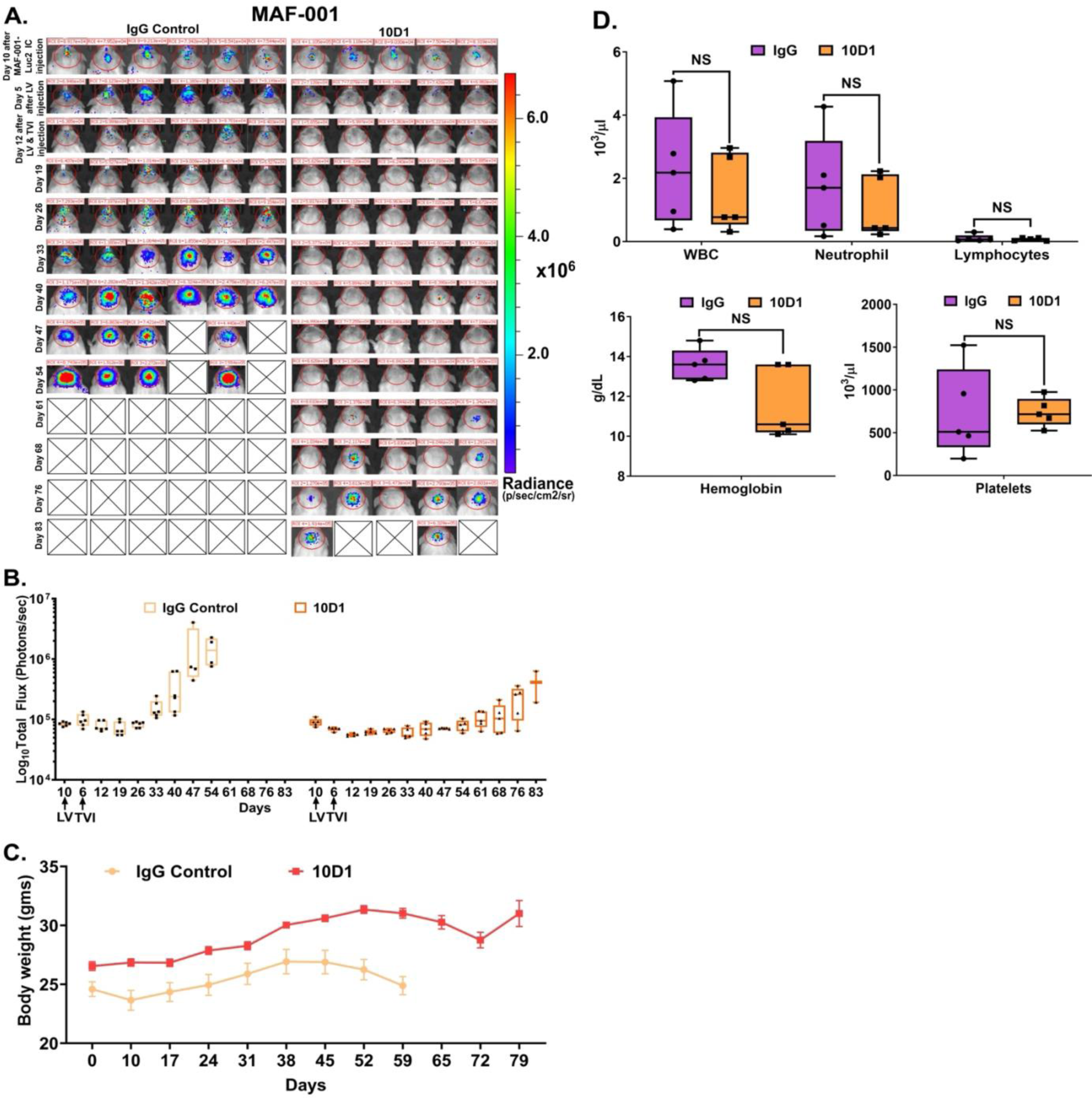
IT injection followed by intravenous injection of 10D1 inhibits tumor growth in MAF-001 xenograft-bearing mice. **A:** MAF-001 PDX model IT followed by IV treatment and its individual BLI of mice in each treatment cohort for different days. **B**: Analysis of BLI for the 10D1 and IgG treatment groups. **C**: Relative body weight of MAF-001 model in both groups. **D**: CBC analysis from the blood collected immediately after animals were euthanized upon reaching their endpoint. Samples were analyzed for WBC (leukocytes), hemoglobin, neutrophils, lymphocytes, and platelets count. Data represented as the mean ± SEM.

**Figure S9:**
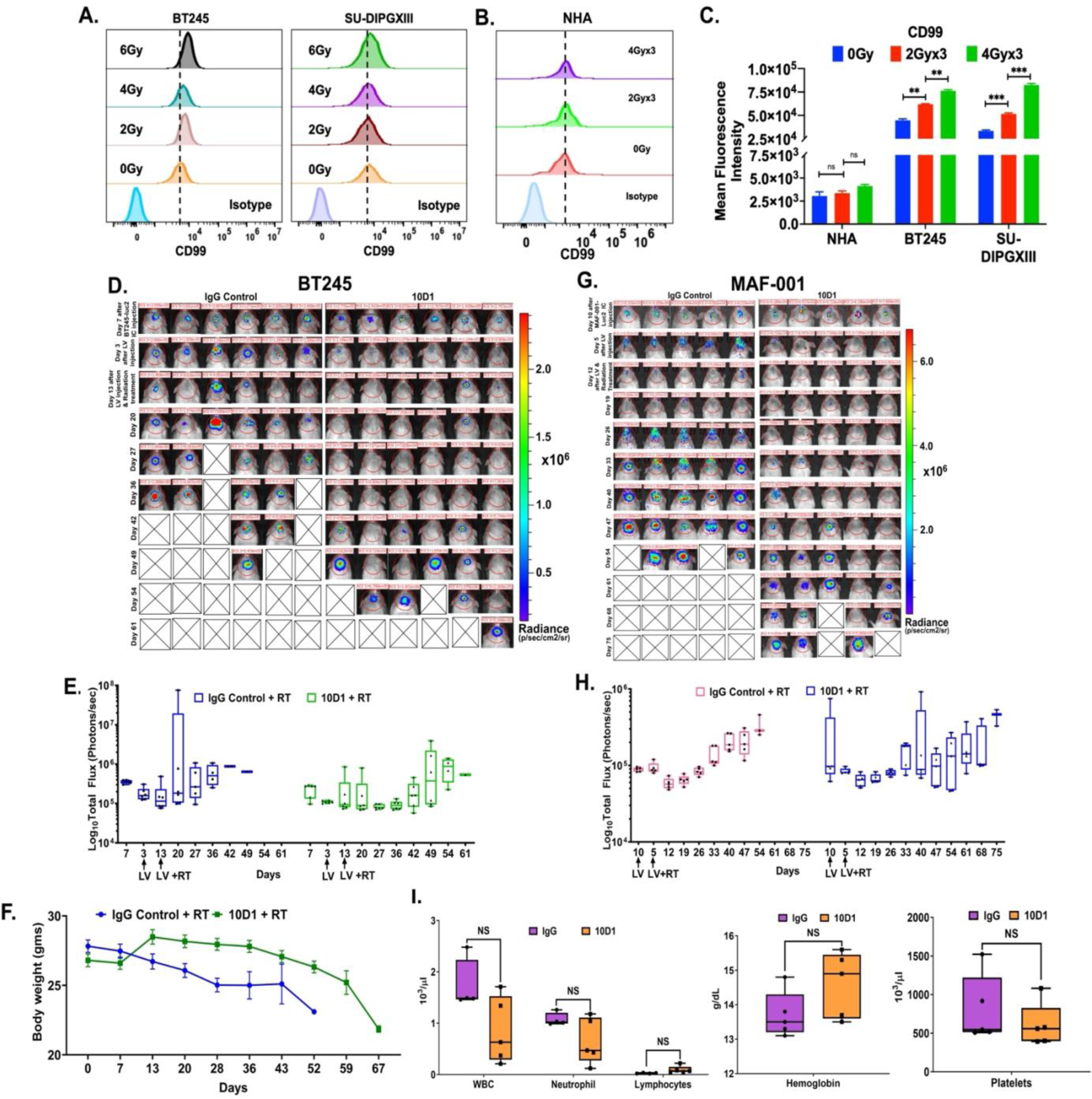
*In vitro* RT study and *in vivo* IT delivery of 10D1 followed by RT inhibits tumor growth and improves survival. **A:** Representative flow cytometry plots showing the expression of CD99 at 24hrs after (a) BT245, and (b) SU-DIPGXIII cells were exposed to a single dose of RT of 2Gy or 4Gy or 6Gy. **B:** Representative flow cytometry plots showing no apparent changes in the expression of CD99 in NHA at 24hrs after they were exposed to FFRT of 2Gy or 4Gy dose for 3 consecutive days. **C:** Quantification of mean fluorescence intensity representing the expression of CD99 after cells were exposed to FFRT with 2Gy or 4Gy dose for 3 consecutive days; (a) NHA from Supplementary Figure S9B, (b) BT245 from Figure 7A(ii) and (c) SU-DIPGXIII from Figure 7A(ii). (ns = not significant, **p<0.01, ***p<0.001) **D**: BT245 and **G:** MAF-001 DMG models and their individual BLI of mice in each treatment cohort for different days. **E & H**: Analysis of BLI for the 10D1 and IgG treatment groups. **F**: Relative body weight of BT245 model in both treatment groups. **I**: CBC analysis from the blood collected immediately after animals were euthanized upon reaching their endpoint. Samples were analyzed for WBC (leukocytes), hemoglobin, neutrophils, lymphocytes, and platelets count. Data represented as the mean ± SEM.

**Figure S10:**
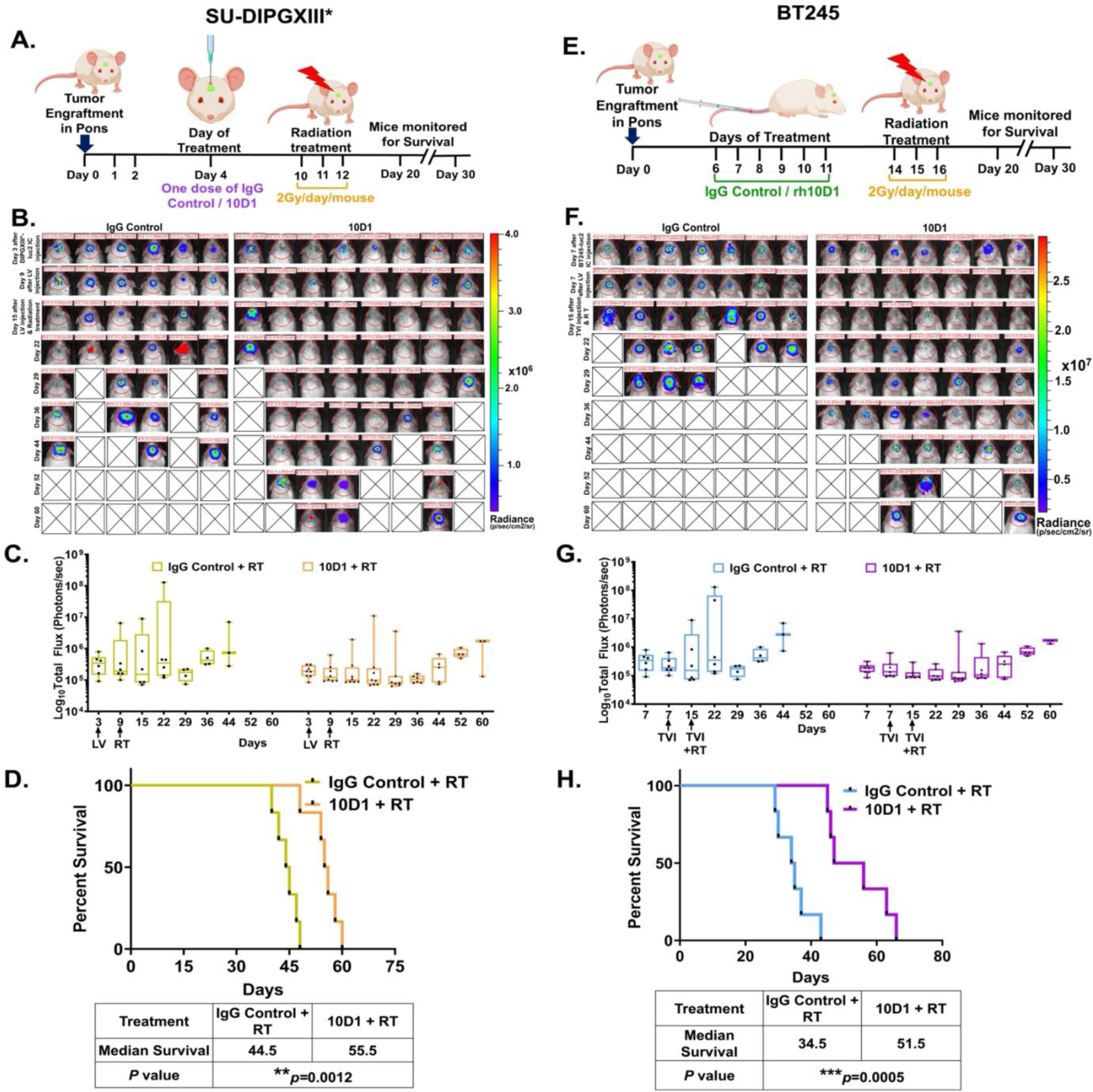
Evaluating different delivery modes of 10D1 in combination with FFRT against DMG xenografts. **A**: Schematic for treatment protocol for SU-DIPGXIII* model. **B**: Representative BLI images of mice after tumor implantation, antibody, and FFRT. **C**: Total Flux values compared with IgG control and treatment groups. Data represented as the mean ± SEM. **D**: Kaplan Meier survival curve shows SU-DIPGXIII* tumor mice treated with one dose of IgG control and 10D1 IT injection followed by FFRT of 2Gy/day for 3 days and compared. Log-rank (Mantel– Cox) test was used to compare groups (**p=0.0012). The table below provides the median survival of the two groups. n = 5 to 6 mice per group. **E**: Schematic for treatment protocol of BT245 model. **F**: Representative BLI of mice after tumor implantation, antibody, and FFRT. **G**: Total Flux values compared with IgG control and treatment groups. Data represented as the mean ± SEM. **H**: Kaplan Meier survival curve shows BT245 tumor mice treated with one dose of IgG control and 10D1 IV injection followed by FFRT of 2Gy/day for 3 days and compared. Log-rank (Mantel–Cox) test was used to compare groups (***p=0.0005). The table below provides the median survival of the two groups. n = 5 to 6 mice per group. Data represented as the mean ± SEM.

## Supplementary Tables

**Table S1: Summary of clinical data, histopathology, DNA methylation classification, and signature genes mutated for patient tumors and cell lines.**

**Table S2: List of antibodies used for ChIP-seq, western blotting, immunohistochemistry, immunofluorescence, and co-immunoprecipitation.**

**Table S3: List of oligonucleotides used for RT-qPCR.**

**Table S4: RNA-Seq data for HSJD-GBM01 (H3.3WT) vs HSJD-GBM01+H3.3K27M (n=2).**

**Table S5: RNA-Seq data for SU-DIPG04 shNull vs shCD99 #1 (n=1).**

**Table S6: Median survival of different tumor models with treatment**

